# Single-cell RNA-seq analysis reveals the crucial role of Collagen Triplex Helix Repeat Containing 1 (CTHRC1) cardiac fibroblasts for ventricular remodeling after myocardial infarction

**DOI:** 10.1101/643130

**Authors:** Adrian Ruiz-Villalba, Juan P. Romero, Silvia C. Hernandez, Amaia Vilas-Zornoza, Nikolaus Fortelny, Laura Castro, Patxi San Martin-Uriz, Erika Lorenzo-Vivas, Paula García-Olloqui, Marcel Palacios, Juan José Gavira, Gorka Bastarrika, Stefan Janssens, Elena Iglesias, Gloria Abizanda, Xabier Martinez de Morentin, Christoph Bock, Diego Alignani, Gema Medal, David Gomez-Cabrero, Igor Prudovsky, Yong-Ri Jin, Sergey Ryzhov, Haifeng Yin, Beatriz Pelacho, Volkhard Lindner, David Lara-Astiaso, Felipe Prósper

**Affiliations:** Program of Regenerative Medicine, Center for Applied Medical Research (CIMA), University of Navarra, Pamplona, Spain; Instituto de Investigación Sanitaria de Navarra (IdiSNA), Pamplona, Spain; Advanced Genomics Laboratory, Program of Hemato-Oncology, CIMA, University of Navarra, Pamplona, Spain; CeMM Research Center for Molecular Medicine of the Austrian Academy of Sciences, Vienna, Austria; Department of Cardiology, Clinica Universidad de Navarra, Pamplona, Spain; Department of Radiology, Clinica Universidad de Navarra, Pamplona, Spain; Department of Cardiovascular Sciences, Clinical Cardiology, KU Leuven, Leuven Belgium; Translational Bioinformatics Unit (TransBio). NavarraBiomed. Pamplona; Department of Laboratory Medicine, Medical University of Vienna, Vienna, Austria; Flow Cytometry Unit, Program of Hemato-Oncology, Center for Applied Medical Research (CIMA), University of Navarra, Pamplona, Spain; Centro de Investigación Biomédica en Red de Cáncer (CIBERONC); Maine Medical Center Research Institute, Scarborough, Maine, USA; Department of Hematology and Cell Therapy, Clinica Universidad de Navarra, Pamplona, Spain

**Author notes:** These authors contributed equally to this work (Adrian Ruiz-Villalba, Juan P. Romero, Silvia C. Hernandez). These authors share senior authorship (Volkhard Lindner, David Lara-Astiaso, Felipe Prósper). Correspondence and requests for materials should be addressed to V.L D.L-A or to F.P.

## Abstract

Cardiac fibroblasts have a central role during the ventricular remodeling process associated with different types of cardiac injury. Recent studies have shown that fibroblasts do not respond homogeneously to heart damage, suggesting that the adult myocardium may contain specialized fibroblast subgroups with specific functions. Due to the limited set of *bona fide* fibroblast markers, a proper characterization of fibroblast population dynamics in response to cardiac damage is still missing. Using single-cell RNA-seq, we identified and characterized a fibroblast subpopulation that emerges in response to myocardial infarction (MI) in a murine model. These activated fibroblasts exhibit a clear pro-fibrotic signature, express high levels of the hormone CTHRC1 and of the immunomodulatory co-receptor CD200 and localize to the injured myocardium. Combining epigenomic profiling with functional assays, we show *Sox9* and the non-canonical TGF-β signaling as important regulators mediating their response to cardiac damage. We show that the absence of CTHRC1, in this activated fibroblast subpopulation, results in pronounced lethality due to ventricular rupture in a mouse model of myocardial infarction. Finally, we find evidence for the existence of similar mechanisms in a pig pre-clinical model of MI and establish a correlation between *CTHRC1* levels and cardiac function after MI.

## INTRODUCTION

Cardiac fibroblasts (CF) represent only 10% of the total number of cells in the myocardium; however, they play a critical role in the structural and mechanical maintenance of the heart ^1,2^. After myocardial injury, provoked by acute ischemia, CF become activated orchestrating a fibrotic response that leads to generation of a collagen scar, which prevents cardiac rupture ^3,4^. The absence of distinct markers for the identification of CF has traditionally hampered our understanding of the fibroblast roles in cardiac homeostasis. However, the generation of fibroblast-specific reporter strains permits to track the origin and roles of fibroblasts in cardiac homeostasis and disease (reviewed in ^3^). Although earlier studies suggested the contribution of endothelial cells and bone marrow cells to activated CF ^3,5,6^, recent studies based on the use of lineage tracing with different Cre drivers indicate that the main source of activated CF are resident fibroblasts that originate from the epicardium during embryonic development ^7–9^. Gene expression profiling studies of different cellular populations after myocardial infarction (MI) have unraveled the pathways mediating CF response to cardiac injury ^10^. However, lineage-tracing studies have also shown that the fibroblast response to cardiac injury is rather heterogeneous, suggesting that different CF subtypes may play a variable role during the healing process that follows MI ^8^. This highlights the need for a better understanding of CF heterogeneity and its impact on processes that mediate repair of the ischemic myocardium

In this study, we used single-cell expression profiling to investigate fibroblast heterogeneity and dynamics during the healing process that follows MI. We identify specific subgroups of CF and characterize the fibroblast subpopulation that becomes activated upon myocardial injury. These responsive fibroblasts present a strong pro-reparative signature, localize to the damaged myocardium, and specifically express high levels of two functionally relevant molecules: the hormone collagen triple helix repeat containing 1 (*Cthrc1*) ^11,12^ and the immunomodulator co-receptor *Cd200*. Using epigenomic profiling we identify the candidate regulatory network that controls their cellular identity, and functionally validate TGFβ/PI3K signaling and *Sox9* as mediators of their specific repair signature. Further, using a knockout model for genetic ablation of *Cthrc1* we provide evidence for the essential role of this specific fibroblast population during cardiac healing. Finally, we highlight the translational relevance of our findings showing cross-species evidence of the contribution of the same CF subpopulation in a pre-clinical swine model of MI and in LV tissue of explanted hearts of end-stage ischemic cardiomyopathy patients.

## RESULTS

### Different fibroblast subpopulations are present in the murine heart

To follow the dynamics of the global cardiac fibroblast population in response to myocardial infarction, we use the *Col1α1-GFP* reporter mouse strain, shown to homogeneously label fibroblasts from different origins ^1,7,13^. To verify the specificity of this model to identify CF, we compared the transcriptional profile of the putative cardiac (GFP^+^/CD31^−^/CD45^−^) and tail fibroblasts, using as external groups, endothelial (GFP^−^/CD31^+^/CD45^−^) and bone marrow-(BM) derived cells (GFP^−^/CD31^−^/CD45^+^) **(Fig. S1a)**. This experiment showed high similarity between the transcriptomes of both sources of fibroblast as compared to endothelial and BM-derived cells **(Fig. S1a-c)**. In line with this, GFP^+^/CD31^−^/CD45^−^ cells isolated from *Col1α1-GFP* cardiac tissue presented a surface membrane profile characteristic of its fibroblastic origin (PDGFRα^+^, mEFSK4^+^/CD11b^−^, CD90^+^/CD45^−^) ^1^ and localized to the site of the injury during the sub-acute and fibrotic phases of cardiac repair at 7, 14 and 30 days post-infarction respectively (dpi) with an increase of GFP^+^ cells after MI (**Fig. 1a** **and Fig. S2a-c**). Taken together, these results indicate that cardiac GFP^+^ cells are *bona fide* fibroblasts, confirming the applicability of the *Col1α1-GFP* model to study CF biology during cardiac repair.

**Figure 1:**
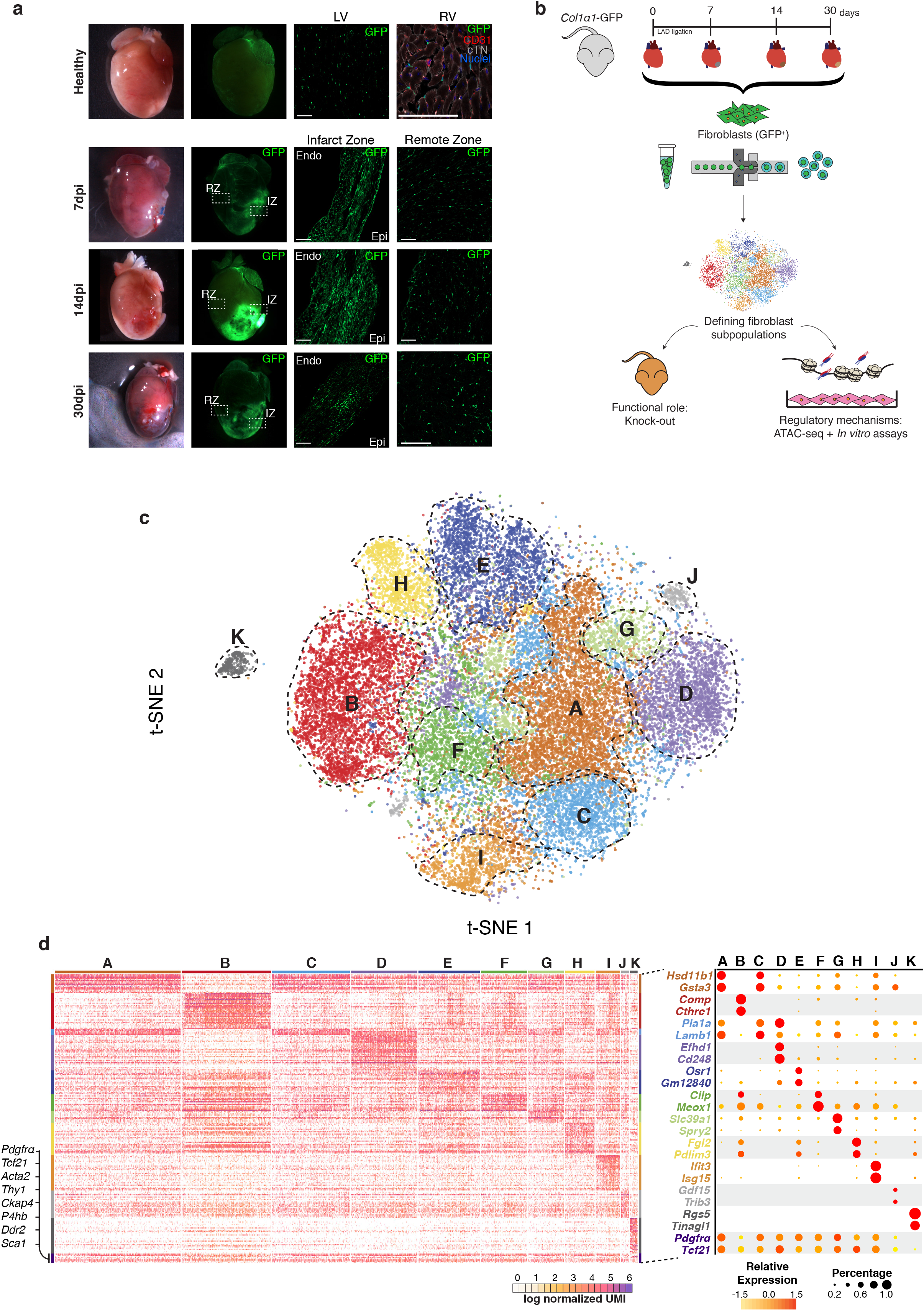
Cardiac fibroblast heterogeneity after myocardial infarction at single-cell resolution. **a)** Overview of representative hearts from healthy myocardium and infarcted myocardium at 7, 14 and 30 dpi using bright field and green fluorescence. Detail of transverse sections showing GFP^+^ cell distribution in the free wall of the left ventricle (LV) in healthy and 7, 14 and 30 dpi in the infarct and remote zones. Detail of GFP^+^ cardiac interstitial cells. Cardiac Troponin-I^+^ cardiomyocytes (cTN) (grey), CD31^+^ endothelial cells (red), nuclei (DAPI, blue). Infarcts were caused by coronary ligation in the left ventricle. Epi, epicardium; Endo, endocardium. **b)** Schematic representation of the experimental design. *Col1α1*-*GFP* mice were subjected to myocardial infarction (MI) by permanent coronary artery ligation and GFP^+^ cardiac fibroblasts (CF) were isolated at different time points (healthy, 7, 14 and 30 dpi). These cells were used for scRNA-seq. Functional role of CF was analyzed with a mouse knock-out model and their regulatory mechanisms with ATAC-seq and *in vitro* assays to validate the hypothesis generated. **c)** t-distributed stochastic neighbor embedding (t-SNE) plot of the global cardiac fibroblast population (29,176 cells) comprising the four different time points (healthy, 7, 14 and 30 dpi). The plot is color coded by the clusters (A-K) identified through unsupervised analysis. Clusters are delimited by dashed lines. **d)** Heatmap showing log normalized UMIs for the analyzed cells using scRNA-seq. Top and side bars indicate clusters. Traditional marker genes for cardiac fibroblasts are indicated (left). Dot plot of expression and specificity of top markers (two) for the identified clusters. Dot size represents percentage of cells per cluster expressing the given marker and color represents the relative expression. *Pdfgra* and *Tcf21* are traditional markers of cardiac fibroblasts.

Next, to characterize at single-cell resolution the fibroblast response after MI, we profiled the single-cell transcriptomes of GFP^+^ CF in healthy myocardium and at 7, 14 and 30 dpi **(Fig. 1b)**. Single-cell transcriptomes from each time point (n = 7,079 cells in healthy, 10,448 7dpi, 8,337 14dpi, and 6,805 30dpi) were subjected to quality control filtering and merged into a single data set of 29,176 single-cell using a canonical correlation approach **(Fig. S3)**. Unsupervised clustering analysis of the global dataset revealed 11 clusters of GFP^+^ cells **(Fig. 1c,d)**. While ten of these clusters (A-J) represent different fibroblast states, characterized by high levels of fibroblast associated molecules (*Pdgfrα*, *Tcf21*, *Acta2*, *Thy1*, *Ckap4*, *P4hb*, *Ddr2*, *Sca1*), cluster K comprises a population with high expression of classical pericyte markers such as *Rgs5*, *Higd1b*, and *Vtn* (**Fig. 1d** **and Fig. S4a,b)**^14^. Of the fibroblast clusters, four of them (B, D, I and J) showed specific expression profiles, representing potential functional CF subgroups, characterized by at least two strong specific markers: *Cthrc1* and *Comp* for cluster B, *Efhd1* and *Cd248* for cluster D, *Ifit3* and *Isg15* for cluster I and *Gdf15* and *Trib3* for cluster J (**Fig. 1c-d**, **and Fig. S4a-c**). The remaining fibroblast clusters (A, C, E, F, G and H) showed less specific transcriptomic identities, which likely reflects intermediate CF cell types within a large fibroblast population (**Fig. 1c-d** **and Fig. S4a-c**).

### Cluster B represents a population of activated fibroblast localized in the infarcted myocardium

To explore the potential role of the different fibroblast subgroups (clusters A-J) during myocardial infarction, we measured their dynamic behavior along the cardiac healing process after MI (healthy and 7, 14 and 30 dpi). While the majority of the fibroblast clusters remained constant in their proportions, only clusters A and B showed a dynamic response to MI. Cluster A comprises 27% of the total fibroblasts in healthy hearts but progressively decreased at 7dpi (25%) and 14dpi (13%), again increasing by 30dpi to a similar level as in healthy myocardium (23%) (**Fig. 2a** **and Fig. S5a**). In contrast, cluster B fibroblasts was almost absent in healthy myocardium (2.3% of the total GFP^+^ cells), rose sharply after MI (12% of total CF at 7dpi and 34% at 14dpi) and began to decrease at 30dpi (12% of the total CF) (**Fig. 2a** **and Fig. S5a**). This pattern suggests that cells in cluster B comprise a transient fibroblast population specific to the healing process. In line with this, cluster B fibroblasts presented an expression signature related to processes involved in tissue repair, such as extracellular matrix organization (ECM), cell proliferation and cell-substrate adhesion (**Fig. 2c** **and Fig. S4d**). This is reflected by high and specific expression of structural molecules including Fibromodulin *(Fmod)* and the Cartilage Oligomeric Matrix Protein (*Comp*) and by enzymes involved in collagen metabolism including Dimethylarginine Dimethylaminohydrolase 1 (*Ddah1*), Lysil oxidase (*Lox),* and Pleiotrophin *(Ptn)* ^15,16^ **(Fig. 2b)**. Periostin (*Postn*), a soluble mediator involved in cardiac remodeling ^9,17,18^, also showed an expression pattern enriched for cluster B fibroblasts, altough its specificity for this fibroblast subpopulation was not as prominent as other repair associated molecules like *Ddah1* and *Lox* **(Fig. S5b and Fig. S6**). On the other hand, *Cthrc1*, a hormone connected to vascular remodeling and fibrotic processes ^19–21^ was identified as a top marker of cluster B fibroblasts at 7 and 14 dpi (**Fig. S5b**). Notably, the pro-repair expression signature of cluster B fibroblasts showed marked dynamics during the healing process as defined by changes in 788 genes (adjusted p-value < 0.05) after MI **(Fig. 2d)**. These changes are organized in three main transcriptional waves: early, intermediate and late, which are enriched in specific functions involved in the repair process. For instance, while collagen related genes and different proteases peak during the intermediate cardiac healing stage (7-14 dpi), several protease inhibitors and cartilage structural genes were upregulated only in the late phase (30dpi), consistent with a balance between synthesis and resorption of ECM components after MI ^22^ **(Fig. S5c)**.

**Figure 2:**
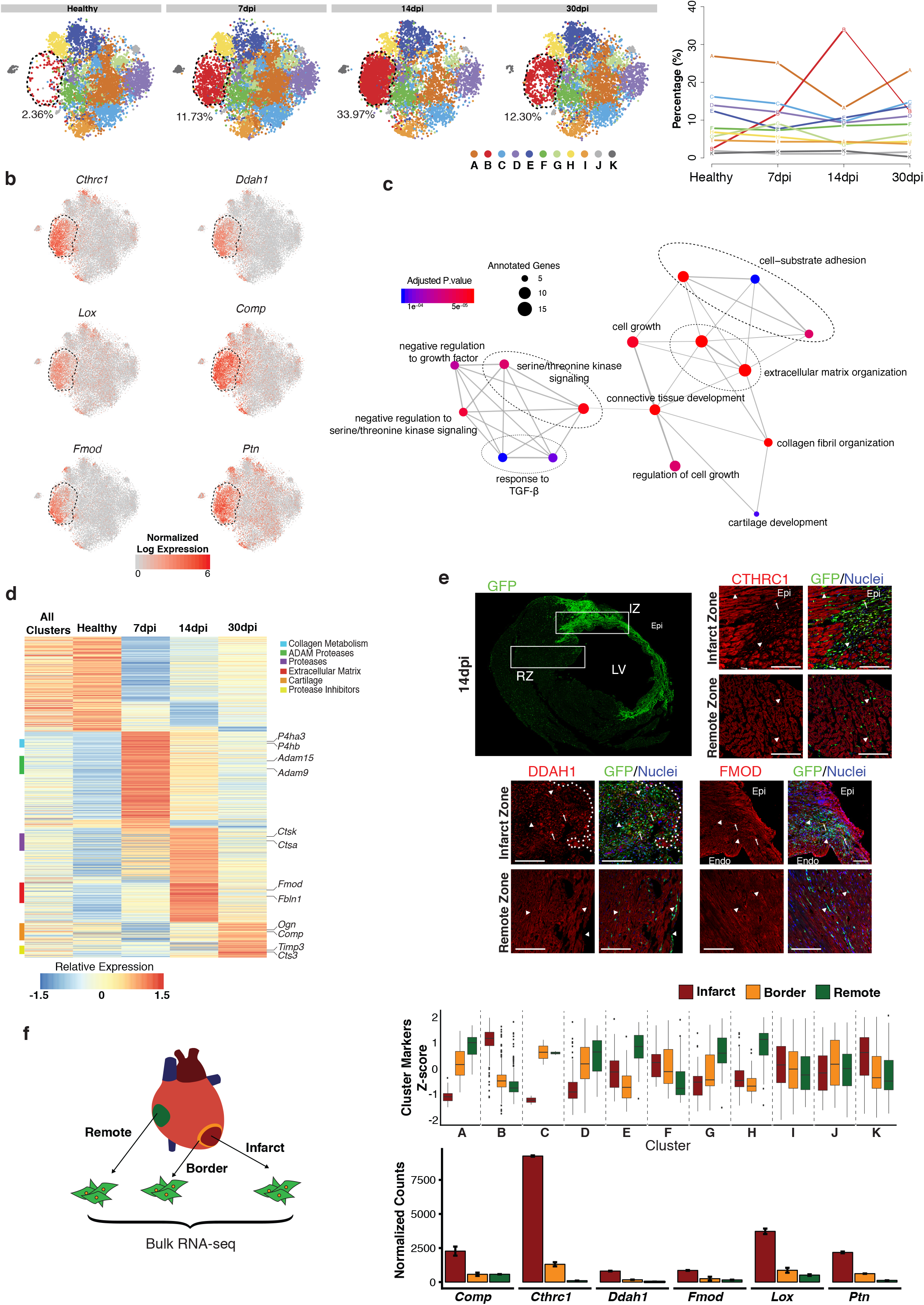
Characterization of cluster B response and tissue location after MI. **a)** (left) t-SNE plot showing cardiac fibroblast heterogeneity in healthy myocardium and at 7, 14 and 30 dpi. Number indicates percentage of cells in cluster B. (right) Plot showing the dynamics of each cluster during the cardiac healing response. For each time point, the proportion of cells belonging to each cluster was calculated and plotted. The lines are color coded according to the identified clusters (A-K) obtained through unsupervised analysis. **b)** Normalized expression of *Cthrc1, Ddah1*, *Lox*, *Comp*, *Fmod* and *Ptn* projected on the t-SNE representation of the global cardiac fibroblast population (29,176 cells) comprising the four different time points (healthy, 7, 14 and 30 dpi). Cluster B is delimited by dashed lines. Rows representing genes with relevant functional categories are labelled with color bars on the left **c)** Network representation of enriched pathways based on IRCFs markers. Dot sizes represent the number of IRCFs markers annotated for each pathway and the color scale represents the statistical significance for each function. **d)** Scaled gene expression heatmap showing transcriptional dynamics of cluster B along the studied time points after MI. **e)** Overview of transverse sections of *Col1α1*-*GFP* infarcted hearts (14dpi). Spatial location of CTHRC1, DDAH1, or FMOD in the infarct zone (IZ) and the remote zone (RZ). GFP^+^ (green, arrowheads), CTHRC1, DDAH1, or FMOD (red), Nuclei (DAPI, blue). Co-localizations are in yellow (arrows). Dotted lines delineate IZ. Epi, Epicardium; Endo, endocardium. **f)** (left) Schematic representation of the zonal gene expression profiling experiment. Cardiac fibroblasts from remote, border and infarct zones were sorted as GFP^+^/CD31^−^/CD45^−^ and their transcriptomes were profiled using low input RNA-seq. (right, above) Boxplot showing the Z-score distribution of cluster markers in the three zonal expression signatures (infarct, border and remote zones) at 7dpi (right, below). Bar plot showing normalized expression values for top cluster B markers in cardiac fibroblasts isolated from infarct, border and remote zones.

The highly specific expression of ECM related genes in cluster B fibroblasts suggests a profound association of this population with the formation of the fibrotic scar. To interrogate the location of these cells in relation to the infarcted myocardium, we performed histological analysis (7, 14, 30 dpi) and zonal transcriptomic profiling of the infarct, border and remote zones at 7dpi. Immunohistochemistry analysis of three specific cluster B markers (*Cthrc1, Ddah1, Fmod*) revealed that CF expressing these molecules are almost exclusively located in the infarct and border zone at 7, 14 and 30 dpi (**Fig. 2e** **and Fig. S7**). In agreement, zonal RNA-seq profiling of the infarcted heart at 7dpi denoted a clear enrichment for cluster B specific expression signature in the infarct area **(Fig. 2f)**. By contrast, none of the remaining clusters (A, C-K) showed a strong connection with the infarct (IZ) or border areas (BZ) at 7dpi, suggesting a looser association of these populations with the damaged tissue. Taken together, our results identify cluster B fibroblasts as a defined subpopulation of CF that (i) emerges in response to myocardial injury, (ii) presents a strong and dynamic pro-fibrotic signature and (iii) localizes to the damaged tissue. These properties suggest that this CF subpopulation plays a major role in the repair process that gives rise to the collagen scar formation, which has led us to call them **Infarct Repair Cardiac Fibroblasts (IRCF)**.

### The immune-modulator co-receptor *Cd200* is a specific marker of Infarct Repair Cardiac Fibroblasts

Next, we searched for specific markers to isolate the IRCF subpopulation. To this purpose, we examined the expression patterns of membrane receptors associated with fibroblast activation in our single-cell dataset. None of the classical activation markers such as *Pgp1* (*Cd44*) ^23^, or fibroblast markers *Thy1* (*Cd90*) or *Pdgfrα* (*Cd140a*) ^24^ showed specificity for IRCFs (cluster B) **(Fig. 3a)**. On the other hand, the immunomodulatory co-receptor *Ox-2*/*Cd200* ^25^ was specifically expressed in cells of clusters B (62.8%) and K (50%), which comprise IRCF and pericytes respectively **(Fig. 3a,b).** Additionally, CD200^+^/GFP^+^ co-localizes with COL1α1 protein to the middle of the scar at 7, 14, and 30 dpi **(Fig. 3d)**. To evaluate whether CD200 could be used to isolate IRCF, we FACS-sorted CD200^−^ and CD200^+^ fibroblasts using CD146 *(Mcam)* to discard pericytes (cluster B 2.9% and cluster K 48.2%) (**Fig. 3c** **and Fig. S8a,b**) and profiled their respective gene expression signatures at 7dpi **(Fig. 3c)**. We found that compared to CD200^−^ fibroblast, CD200^+^ CF expressed much higher levels of cluster B markers and display a significant similarity with the aggregated expression signature of cluster B cells (**Fig. 3c** **and Fig. S8c**). This confirms the specific presence of the co-receptor CD200 in the membrane of IRCF (cluster B CF), highlighting an alternative to isolate activated fibroblasts after MI.

**Figure 3:**
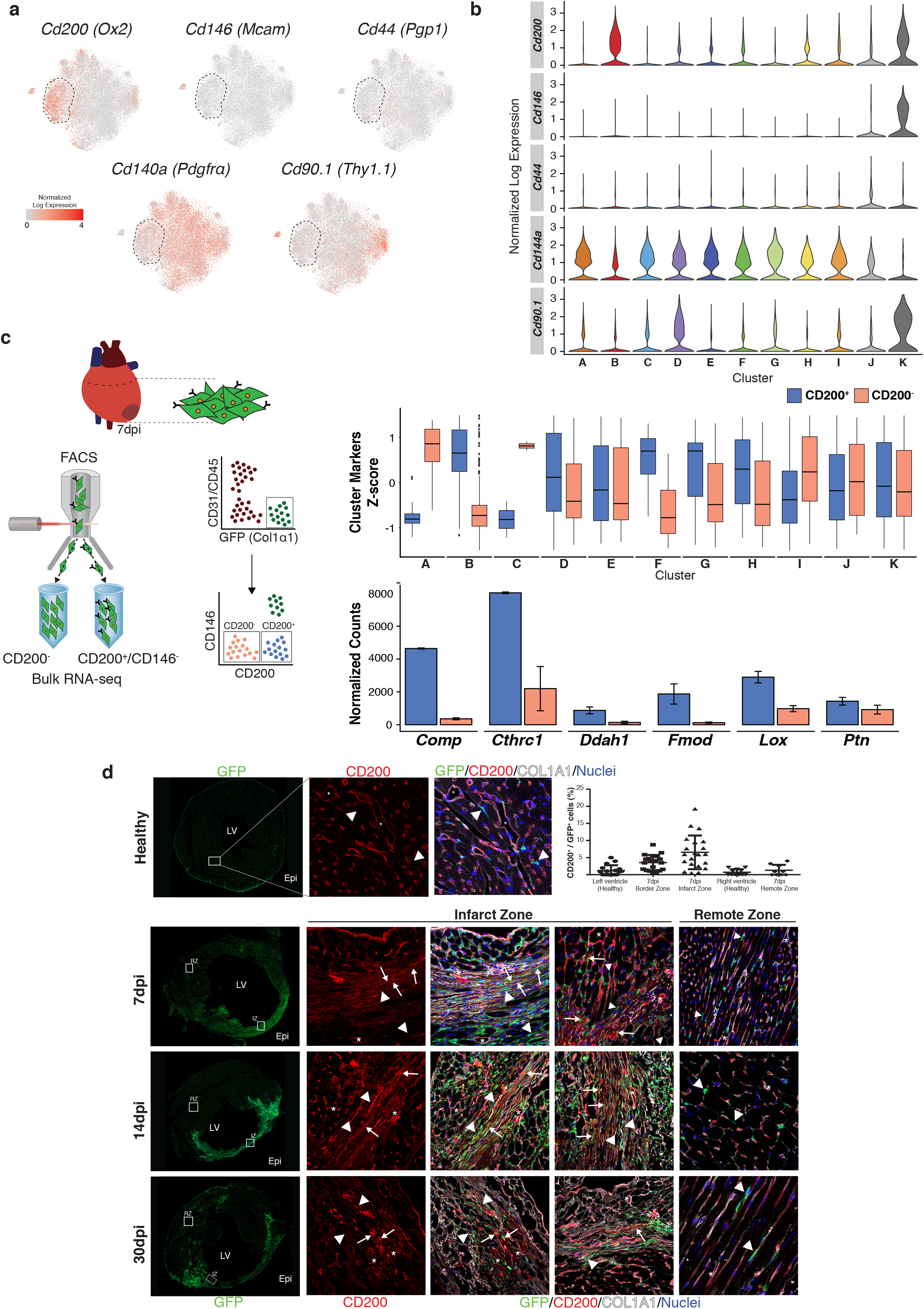
CD200 is the most specific cell surface marker for IRCF. **a)** t-SNE representation showing single cell normalized expression of selected surface markers in the pooled cardiac fibroblast population that comprises the four different time points (healthy, 7, 14 and 30 dpi). Cluster B is delineated by dashed lines. **b)** Violin plots showing the expression of *Cd200 (Ox2), Cd146 (Mcam), Cd44 (Pgp1), Cd140a (Pdgfrα),* and *Cd90.1 (Thy1.1)* in every cardiac fibroblast cluster. **c)** (left) Schematic representation of the strategy employed to isolate CD200^+^ cardiac fibroblasts (GFP^+^/CD200^+^/CD146^−^) and CD200^−^ cardiac fibroblasts (GFP^+^/CD200^−^/CD146^−^) at 7dpi; CD146 was used to discard the CD200^+^ pericyte population (right, above). Boxplot representation of Z-score distributions for cluster markers in CD200^+^ and CD200^−^ cardiac fibroblasts (above). (below) Normalized expression bar plots for top IRCF markers in both CD200^+^ and CD200^−^ cardiac fibroblasts. **d)** Representative transverse sections of *Col1α1-GFP* hearts at different time points. Immunofluorescence analysis of GFP^+^ (green), CD200^+^ (red), Collagen1α1^+^ (grey) and DAPI/nuclei (blue) in healthy left ventricle and infarcted (IZ) (left) and remote zones (RZ) (right) at 7, 14 and 30 dpi. Co-localization of GFP^+^ and CD200^+^ in yellow (arrows) in IZ at different time points after MI (and light yellow where both co-localize with Collagen1α1). Arrowheads indicate GFP^+^/CD200^−^ cells and asterisks indicate GFP^−^/CD200^+^. Quantification of GFP^+^/CD200^+^ cells in healthy and at 7dpi in different regions of the heart (right, top). LV, left ventricle; Epi, epicardium; IZ, infarct zone; RZ, remote zone.

### Potential transcription factors regulating IRCF response

To explore the regulatory mechanisms involved in the activation of cluster B fibroblasts (IRCF), we profiled the chromatin accessibility patterns (ATAC-seq) of CD200^+^ and CD200^−^ fibroblasts at 7dpi, and of global CF population (GFP^+^) in healthy myocardium and at 7, 14 and 30 dpi **(Fig. 4a)**. We observed the highest chromatin accessibility for cluster B specific loci in CD200^+^ CF (**Fig. 4a** **and Fig. S8c**). Next, to identify which transcription factors (TFs) regulate IRCF identity, we performed motif analysis on IRCF specific distal accessible regions (>1.5 Kb away from TSS). This analysis identified a group of potential regulators comprising: members of the AP-1(JUN) and NF-κB families (RELA), which are involved in inflammatory responses; SMAD factors, known to mediate fibrotic processes ^26^; TEAD factors, known regulators of fibroblast fate; and *Runx1* a central regulator in relevant developmental processes **(Fig. 4b)**. In addition, to circumvent limitations due to biased motif annotations, we leveraged public ChIP-seq databases to find TFs whose binding patterns are enriched in the vicinity of cluster B genes. This approach confirmed many of the candidates identified in the motif approach and highlighted additional TFs such as SOX9, RXRA whose binding footprints were not significantly enriched in our motif analysis **(Fig. 4c)**. Most of the uncovered potential regulators, such as AP1 and SMAD factors, are ubiquitous factors involved in mediating fast signal transduction responses and, not surprisingly, their single-cell expression patterns showed no specificity for cluster B fibroblasts (IRCF) **(Fig. S9a)**. On the other hand, *Runx1* and *Sox9* belong to known lineage regulatory TF families and, interestingly, their single-cell expression patterns showed specificity for cluster B CF, suggesting a central role in regulating the cellular identity of IRCF **(Fig. 4b,c)**. To validate this hypothesis, we overexpressed *Sox9* and *Runx1* in cultured CF obtained from healthy hearts and then measured their ability to induce a cluster B specific signature using RNA-seq (**Fig. 4d** **and Fig. S9b**). With the exception of *Cthrc1 and Ptn*, overexpression of *Runx1* failed to induce specific transcripts of cluster B fibroblasts. However, *Sox9* overexpression induced the expression of 23% of cluster B specific genes (28 genes FC>1.5 p-value <0.05), indicating the potential of *Sox9* to promote an activated IRCF phenotype (**Fig. 4d** **and Fig. S9b**).

**Figure 4:**
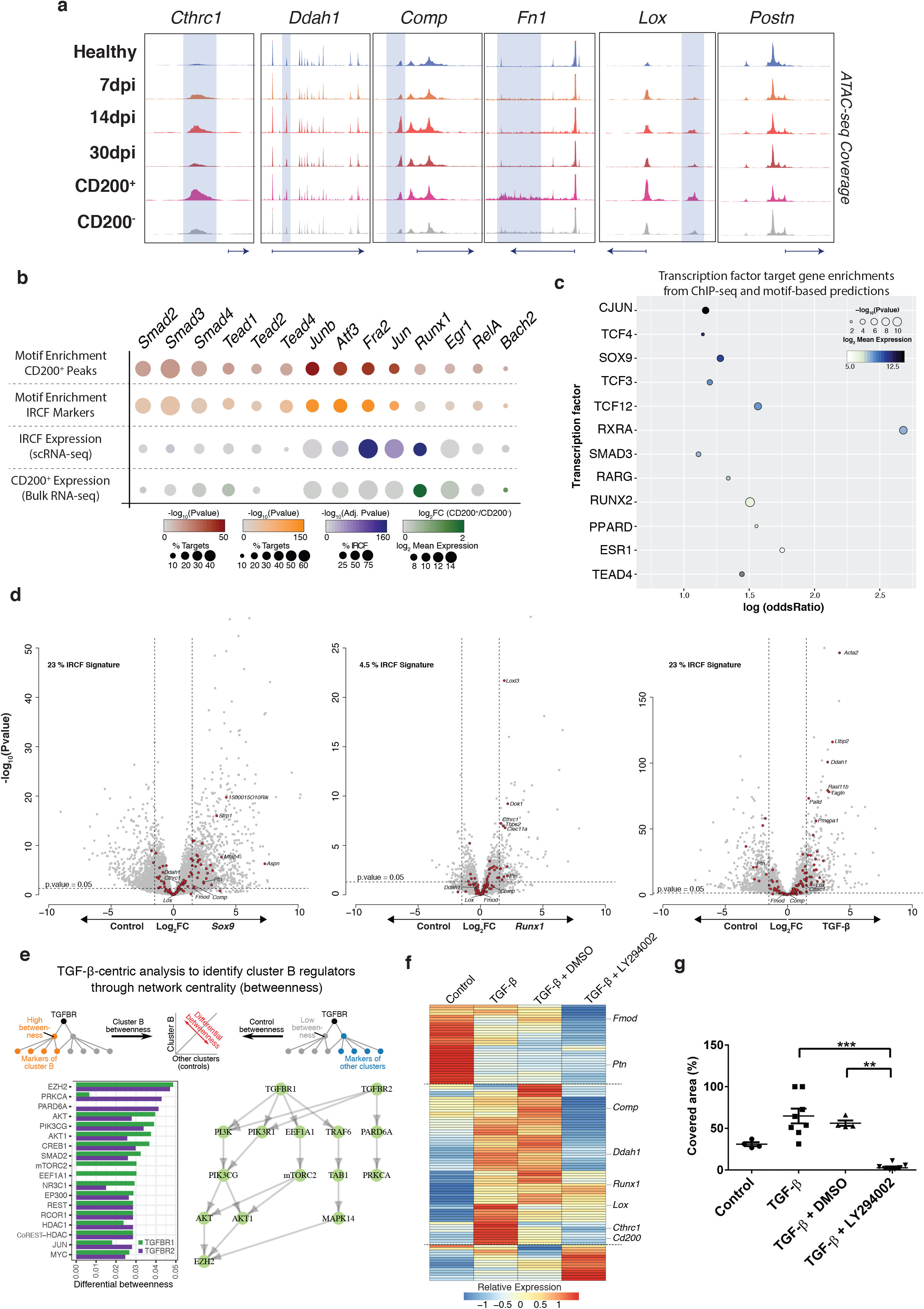
Regulation of infarct repair cardiac fibroblast. **a)** Genome browser snapshots showing the accessibility profiles of representative loci in the global cardiac fibroblast population at 0 (healthy), 7, 14 and 30 dpi and in the CD200^+^ and CD200^−^ cardiac fibroblast subpopulations isolated at 7dpi. Shadowed areas mark distal regulatory elements displaying increased accessibility in the CD200^+^ subpopulation. **b)** Dot plot representing motif enrichment and expression specificity of the potential transcription factors mediating the response of IRCFs (cluster B/CD200^+^ CF). The first row represents the values obtained from the motif analysis of the CD200^+^ specific accessible distal regulatory elements (+/− 1.5 Kb from TSS). The second row represents the values obtained from the motif analysis of the distal regulatory elements found within cluster B specific loci. Third and fourth rows show the expression values of the transcription factors in scRNA-seq and CD200^+^ bulk RNA-seq datasets, respectively. **c)** Transcription factor target gene enrichments from different ChIP-seq experiments and motif predictions. The x-axis denotes the odds ratio of the transcription factor enrichment. Dot size represents the enrichment p-value and color represents the log2 transformed expression in CD200^+^ bulk RNA-seq. **d)** Volcano plots showing differential gene expression of ex vivo grown CF overexpressing *Sox9* (left), *Runx1* (middle) and treated with TGF-β (left). Genes with log Fold Change of +/− 1.5 and p-value <0.05 were considered differentially expressed. Red dots represent IRCF markers. **e)** TGF-β network centrality analysis revealed the non-canonical PI3K-AKT pathway to be related with IRCF markers. **f)** Heatmap showing the relative expression of IRCFs markers in non-treated (Control), TGF-β, TGF-β + Vehicle, and TGF-β + LY294002 cultured CFs. **g)** Quantification of area covered by cultured CFs after 23h of treatment in wound healing experiment in CF that were untreated, or treated with TGF-β, TGF-β + DMSO, or TGF-β + LY294002.

### IRCF response is partially mediated by non-canonical TGF-β signaling

TGF-β signaling has been involved in a variety of fibrotic processes ^26^. In our data, gene set enrichment and motif enrichment analyses of cluster B specific genes identified TGF-β and SMADs, pointing at canonical TGF-β signaling as a key pathway mediating IRCF response (**Fig. 2c** **and** **Fig. 4b**). To further dissect the signaling route connecting TGF-β stimulation to IRCF specific genes, we build a network spanning protein signaling and gene-regulatory interactions and used a graph-theory-based analysis on our single-cell expression dataset to identify the signaling proteins that are “in between” TGF-β receptors and cluster B marker genes. Surprisingly, rather than the canonical/SMAD pathway, this analysis highlighted the non-canonical TGF-β/PI3K-Akt signal transduction branch as the main driver of cluster B gene expression (**Fig. 4e**). To validate the implication of TGF-β/PI3K-Akt signaling pathway in mediating IRCF response to MI, we stimulated CF *in vitro* with TGF-β with and without LY294002, a known PI3K-Akt inhibitor **(Fig. S9c)** ^27^. TGF-β stimulation recapitulated 25% of the cluster B specific expression pattern (33 genes FC>1.5 p-value<0.05) confirming the potential of TGF-β to trigger an IRCF response **(Fig. 4d)**. However, this effect was significantly abolished upon PI3K-Akt inhibition (**Fig. 4f** **and Fig. S9d**). In addition, using a wound healing model, we observed that the ability of fibroblast to induce migration was enhanced by TGF-β but abolished in the presence of the PI3K inhibitor (**Fig. 4g** **and Supplementary Movie 1**). These results emphasize the relevance of the non-canonical TGF-β1/PI3K-Akt signaling in controlling cluster B specific expression. This results do not rule out a potential role of the canonical TGF-β signaling as high SMAD motif enrichment was identified in accessible chromatin of cluster B cells. Moreover, the absence of SMAD proteins in our betweenness analysis might be due to the constitutive expression of these factors across the single-cell fibroblast clusters. Thus, our results suggests that TGF-β signaling mediates IRCF fibrotic response during MI via both, canonical/SMAD and non-canonical pathways.

### CTHRC1 mediated IRCF activity is essential for the formation of the healing scar

Having explored molecular mechanisms that regulate IRCF identity, we set out to assess their physiological roles during cardiac repair. To this end, we studied the progression of cardiac repair in mice with genetic ablation of the hormone *Cthrc1* (*Cthrc1-KO*) ^21^, an IRCF top marker that has been linked to several fibrotic processes ^19^. *Cthrc1-KO* mice undergoing MI showed a dramatic decrease in survival due to rupture of the free wall of the left ventricle between 5-7 dpi in comparison with wild-type (WT) animals (70% KO *vs* 20% WT) **(Fig. 5a)**. This was reflected by the reduction in the fractional shortening (FS) (p=0.011 in comparison with WT) and an increment in LVIDd and LVIDs observed at 7dpi (p=0.013 and 0.009 respectively in comparison with WT) **(Fig. 5b)**. These results indicate a crucial role for CTHRC1 in myocardial repair. Of note, CTHRC1 expression was not detected in non-fibroblastic cardiac cells (endothelial, BM-derived or cardiomyocytes) in infarcted hearts of WT animals (**Fig. 5c** **and Fig. S10a,b,d**). This highlights the specificity of *Cthrc1* for fibroblasts and indicates that the dramatic phenotype observed in *Cthrc1-KO* animals is due to the ablation of this gene in CF.

**Figure 5:**
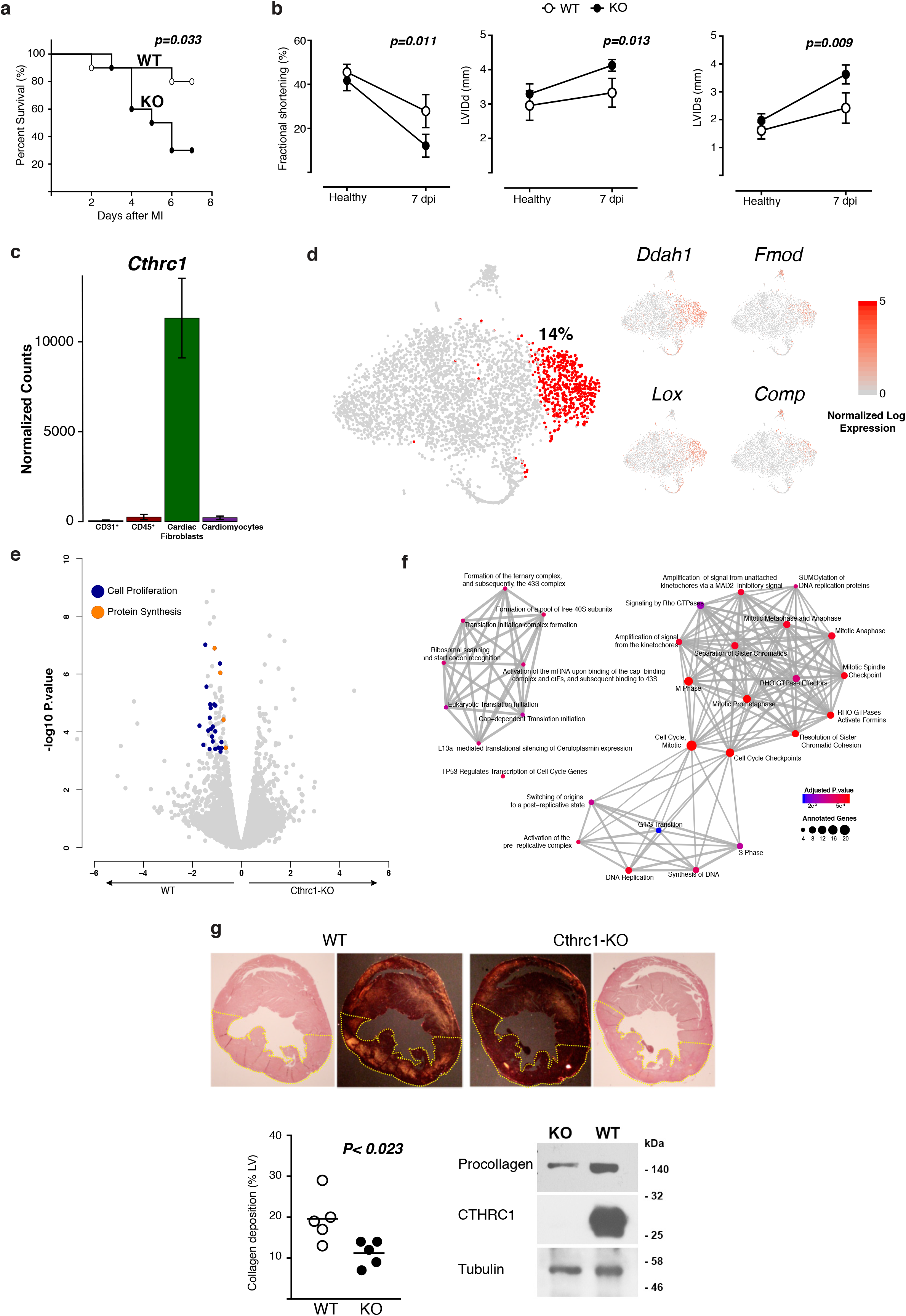
*In vivo* effect of Cthrc1 deletion after myocardial infarction. **a)** Kaplan-Meier survival curves after MI in wild type (WT, n=10) and *Cthrc1* knockout (KO, n=10) mice. Survival analysis was performed by *log*-rank test. **b)** Cardiac function in *Cthrc1*-KO that survived MI (closed circle) compared to WT (open circle) animals using echocardiographic examination before and at 7dpi. *From left to right:* Left ventricular fractional shortening (Fractional shortening (FS, %)), End-diastolic left ventricular internal diameter (LVIDd), End-systolic left ventricular internal diameter (LVIDs). n = 10 for WT and KO before MI, n=8 (WT) and n=3 (KO) at 7dpi. p-value was calculated using an unpaired *t*-test. **c)** Normalized expression bar plots of *Cthrc1* in endothelial cardiac cells (CD31^+^), bone marrow derived cardiac cells (CD45^+^), cardiac fibroblasts (mEFSK4^+^/CD31^−^/CD45^−^) and cardiomyocytes obtained from wildtype hearts at 5dpi. **d)** t-SNE representation of 4,189 cardiac fibroblasts (mEFSK4^+^/CD31^−^/CD45^−^) obtained from one Cthrc1^−/−^ heart at 7dpi. The number represents the percentage of cluster B like fibroblasts present in the Cthrc1^−/−^ heart at 7dpi. t-SNE representation for IRCF cluster markers: *Ddah1, Fmod, Lox, Comp* in cardiac fibroblasts isolated from Cthrc1^−/−^ hearts at 7dpi. **e)** Volcano plots showing differential gene expression between wildtype and *Cthrc1*^−/−^ cardiac fibroblasts at 5dpi. In this case cardiac fibroblasts from wildtype and control were sorted as mEFSK4^+^/CD31^−^/CD45 **f)** Network representation of enriched gene-ontology categories for genes down regulated in the *Cthrc1*^−/−^ cardiac fibroblasts. **g)** Representative image of collagen deposition in the left ventricle of wildtype (left) and *Cthrc1*^−/−^ (right) mice viewed under polarized light of picrosirius red stained sections (above). Quantification of collagen deposition in the left ventricle in both genotypes at 3dpi analyzed by unpaired *t*-test (Wildtype, open circle; *Cthrc1^−/−^*, closed circle) (left, below). Western blot probed for Procollagen-I, CTHRC1 and tubulin of lysates from cultured adult CF derived from *Cthrc1*^−/−^ and wildtype mice (right, below).

To unravel the mechanisms underlying *Cthrc1* beneficial effects during MI, we analyzed the bulk and single-cell transcriptomes of CF in the *Cthrc1-KO* background between 5-7 dpi. Surprisingly, we found that the proportion of cluster B-like cells in *Cthrc1-KO* hearts is similar to the proportion in WT mice at 7dpi, indicating that the absence of *Cthrc1* does not affect the induction of cluster B CF after MI (**Fig. 5d** **and Fig. S10e**). However, even though the emergence of cluster B fibroblasts was not affected, differential expression analysis between WT and KO strains at 5dpi detected significant differences between WT and KO CF populations (113 genes adjusted p-value <0.05) **(Fig. 5e)**. In particular, we found that KO CF failed to activate genes related to cell division, proliferation, and protein synthesis upon MI (**Fig. 5e,f**). This indicates that *Cthrc1* mediates its cardioprotective effect by activating proliferation and protein synthesis in CF, functions that are crucial for the formation of the fibrotic scar. In line with this, we found that collagen deposition in KO infarcted hearts was significantly decreased (50%), reflecting a compromised protein synthesis capability in CF upon CTHRC1 disruption (**Fig. 5g** **and Fig. S10c**). Taken together, these results suggest that IRCF orchestrate cardiac repair at two levels: by deposition of ECM molecules and by secreting molecules, such as CTHRC1, that reinforce the proliferative and ECM deposition abilities of CF.

### An IRCF like expression signature is detected in pre-clinical models of myocardial infarction

To assess the translational potential of our findings, we examined whether the IRCF expression signature can be detected in a pre-clinical pig model of MI (ischemia/reperfusion (I/R)). To this aim, twenty-three pigs were subjected to MI and sacrificed at 8 or 180 dpi for gene expression profiling (RNA-seq) and histological analysis **(Fig. 6a)**. In addition, 2 healthy pigs were used as controls. Since fibroblast surface markers are poorly characterized in porcine models, we analyzed the global myocardial transcriptome of total cell lysates obtained from infarct (IZ) and remote zones (RZ) **(Fig. 6b)**. Despite the lack of cell type resolution, our approach was able to detect the induction of 33% and 32% of the IRCF signature at 8dpi and 180dpi respectively in the IZ but not in RZ **(Fig. 6b)**. In agreement, CTHRC1^+^ cells were spatially located to the IZ but not in the RZ, in the same region where deposits of COL1α1 and POSTN were found **(Fig. 6c)**. These results indicated that a population of CF similar to the murine IRCF appears after MI in our swine pre-clinical model.

**Figure 6:**
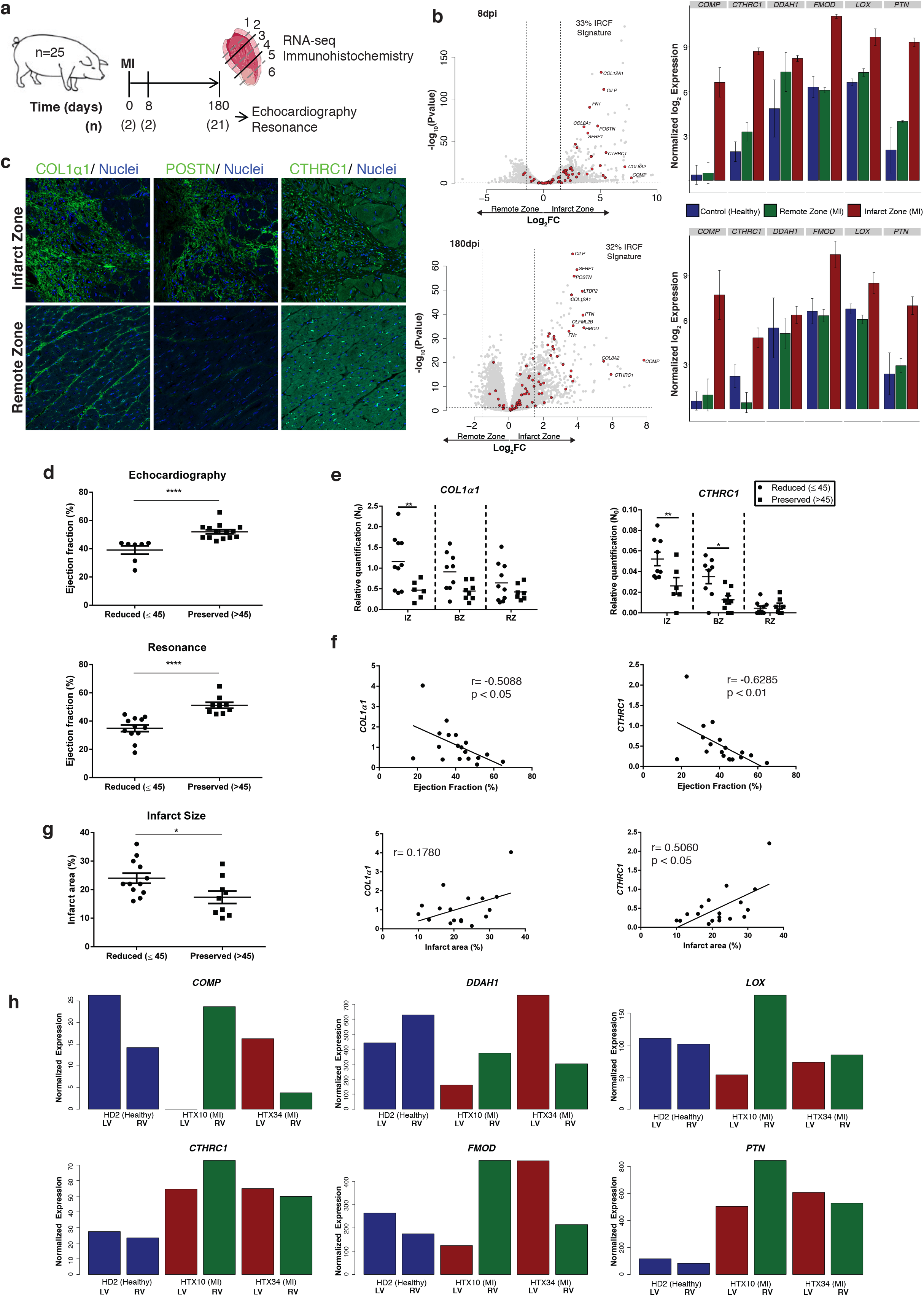
CTHRC1 expression and functional correlation in a preclinical model of MI. **a)** Schematic representation of the experimental design. Twenty-five pigs were included in the study, twenty-three were submitted to Ischemia/Reperfusion (I/R) surgery and 2 animals were used as controls. Cardiac function was determined in 21 pigs 6 months after the surgery using echocardiography and MRI. **b)** Zonal transcriptomic profiling of swine models at 8dpi (top) and 180dpi (bottom). Volcano plots (left) show differentially expressed genes between infarct and remote areas. Red dots indicate porcine transcripts homologous to IRCF markers. Differential gene expression analysis detects 33% and 32% of the IRCF signature overexpressed in the infarct zone at 8dpi and 180dpi respectively. Normalized expression bar plots (right) for top IRCF markers at 8dpi and 180dpi in the remote (green) and infarcted (red) heart areas. **c)** Immunohistochemistry of COL1α1, POSTN and CTHRC1 in representative areas of infarct and remote zones at 8dpi. **d)** Distribution of pigs with reduced (≤45%, circles) or preserved (>45%, squares) ejection fraction (EF) at 180dpi measure by echocardiography or MRI. **e)** Expression levels of *COL1a1* and *CTHRC1* in different anatomical regions of the infarcted heart between pigs with reduced EF (circles) and preserved EF (squares). **f)** Correlation between EF and the level of expression of *COL1a1* and *CTHRC1* in the infarct zone. **g)** Quantification of infarct area (+) using MRI. Correlations between the infarct area and the level of expression of *COL1a1* and *CTHRC1* in the infarct zone. **h)**Normalized expression bar plots for top IRCF markers in zonal biopsies of 2 patients (HTX10 and HTX34) and a control (HD2). EF, ejection fraction; IZ, infarct zone; BZ, border zone; RZ, remote zone. LV, left ventricle; RV, right ventricle * p=0.05, ** p=0.01, *** p=0.0001

The severity of the Cthrc1^−/−^ phenotype suggests that the levels of *Cthrc1* could be an important factor influencing the outcome of the cardiac healing process after acute ischemic injury. To assess this possibility we measured the expression of *CTHRC1* in animals with preserved (>45%) and reduced (≤45%) Ejection Fraction (EF) at 180 dpi **(Fig 6d)**. Surprisingly, even though in the early stages of cardiac healing in mice CTHRC1 is critical to prevent cardiac wall rupture, we found that at later stages (180 dpi) the level of *CTHRC1* expression in the wounded myocardium (BZ, IZ) is significantly higher in the of group of pigs with cardiac dysfunction. Accordingly, a negative correlation between the level of expression of *CTHRC1* and the EF in the IZ was found (p<0.01 and p<0.05, respectively) **(Fig. 6e,f)**. On the other hand, a positive correlation between infarct area and the expression of *CTHRC1* was found **(Fig. 6g)**. These results suggest that levels of *CTHRC1* are associated with the fibrotic response. Similarly, levels of *COL1α1* were also associated with cardiac dysfunction **(Fig 6e)**. Finally, gene expression analysis in samples from left and right ventricle (LV and RV) in 2 patients with MI revealed an increase in *CTHRC1* levels and an IRCF like signature in human heart biopsies derived from the ischemic territory of patients with ischemic cardiomyopathy but not in the remote, non-ischemic zones **(Fig. 6h)**. These findings highlight the potential role of *CTHRC1*^+^ IRCF in orchestrating the cardiac repair process in patients after MI and suggest that cardiac levels of CTHRC1 may be a biomarker of cardiac dysfunction after MI

## DISCUSSION

As the main cellular component responsible for cardiac healing after MI, CF are an attractive therapeutic target. However, the lack of *bona fide* markers that define CF, the evidence of their heterogeneity and the limited understanding of the molecular mechanisms underlying their activation has precluded the development of successful therapies targeting fibrotic remodeling. Using a reporter mouse strain that unequivocally identifies CF, we have contributed not only to characterize the unappreciated heterogeneity of the CF population and its response to MI but more importantly, we identify a specific population of activated CF responsible for the response to the ischemic injury (IRCF), establish specific markers, demonstrate their functional role after MI and provide insights into their mechanism of regulation.

Non-myocyte cardiac cell heterogeneity has been recently characterized based on FACS sorting different populations of immune, endothelial and cardiac fibroblast ^14,28,29^ or on the usage of lineage tracing models such as PDGFRα ^30^ or POSTN ^9^ reporter mice. Most of these studies, focus on the early phase after cardiac injury ^30^ and assessed other pathological conditions ^9,28,29^. In contrast, we focused on the heterogeneity of CF, analyzing a significantly larger number of cardiac fibroblasts collected from healthy and infarcted mice (29,176). In addition, our cell selection strategy was based on a different tracing model, potentially explaining some of the differences observed. For instance, Kretzschmar *et al.* identified 11 clusters within 282 CF at 14dpi using Mki67-RFP reporter mice ^28^. In this study, the authors described a subpopulation of activated CF located in the scar region with a similar function that IRCFs. However, both populations of CF are characterized by different markers, most probably because of the low number of Mki67^+^ CF included in the scRNA-seq study, and the reporter mice used. Interestingly, Faberhi *et al.* described 9 subpopulations of PDGFRα^+^ CF with a similar transcriptomic profiles that our subpopulations of *Col1α1-GFP*^+^ CF at 7dpi ^30^. However, our data indicate that CF that express *Pdgfrα* are contained within *Col1α1-GFP*^+^ CF (**Fig. 3a** **and Fig. S2a**), suggesting that PDGFRα-GFP CF are included into the IRCF population.

A recent study has defined the CF response after MI using three different mouse models of cell lineage tracing ^31^. The results of this study, based on stage specific transcriptomics, suggest a model of CF kinetics that includes the initial proliferation followed by activation and migration of CF and eventually the conversion of these activated CF into non-proliferating, matrix-producing fibroblast named “matrifibrocytes” ^31^. IRCF show a similar transcriptional signature and anatomical location into the fibrotic scar to “matrifibrocytes” (**Fig. 2e** **and Fig. S5b**). Here, we provide additional insights into the functional role and transcriptional regulation demonstrating that IRCF are specifically located in the infarct zone but not in the remote areas of the myocardium as shown both by zonal RNA-seq as well as immunohistochemistry analysis. In addition, this population is almost absent in healthy myocardium but accounts for more than 30% of the CF at 14dpi, further supporting their role in the healing process and making IRCF an attractive target for therapy. Surprisingly, although deletion of *Cthrc1* was associated with increased mortality due to the reduction of the fibrotic response, the specific population of IRCF was still detected in the myocardium using scRNA-seq and immunohistochemistry, even though significant changes in their proliferative signature were observed. Future studies should addressed whether deletion of IRCF using Cre-inducible DTR transgenic mice models instead of deletion of the *Cthrc1* hormone would result in a different outcome ^32^. Based on their proliferation, a population of activated CF identified by their expression of *Fstl1* and responsible for preventing cardiac rupture after MI has recently been characterized ^28^ which could also be part of our IRCF.

CTHRC1 is a secreted protein that has the ability to inhibit collagen matrix synthesis through inhibition of TGF-β signaling ^20,21,33^ and that appears as a top marker gene for IRCF. Although expression of *Cthrc1* has been previously identified as a potential marker for activated CF after MI in mice ^9,30^ and humans ^29^, the longer follow-up in our study (up to 30dpi) may explain why it was not previously identified as a key player in the cardiac repair process. According to our results, expression of *Cthrc1* peaks at 14dpi and decreases thereafter. Besides the potential for defining the population of IRCF, this dynamic suggest that expression of *Cthrc1* may be responsible for promoting the fibrotic response after MI, an essential part of the healing process ^3,34^. Targeted ablation of other markers of activated CF such as *Postn* ^9,35^ or *Fstl1* ^28^ have been associated with increased mortality after MI. Interestingly, timing of *Postn* deletion seems to modify the outcome after MI ^9,35^. Our results in a pre-clinical model of MI provide additional translational relevance to our findings as expression of *CTHRC1* was upregulated in the infarct zones with identification of CTHRC1 expressing cells in these zones and not in the remote myocardium, consistent with findings in our mice models. Unlike studies performed in mice, RNA-seq studies in pig and patients were performed on whole cell lysate from infarct tissue. Expression of *CTHRC1* has been mostly detected in cardiac fibroblasts in the heart as well as other areas outside the heart tissues such as brain ^36^ and bone ^12^, with bone being the source of CTHRC1 detectable in circulation ^37^. The relation between *CTHRC1* expression and function indicates that CTHRC1 may represent an interesting biomarker of cardiac remodeling after MI.

Expression of *Cd200* was detected in more than 60% of the IRCF but more importantly, was not present in other CF populations. CD200, previously known as OX2, is a membrane glycoprotein initially described to be broadly distributed in activated immune and endothelial cells that plays an important role in immunosuppression and regulation of anti-tumor activity ^38–40^. A variety of *in vitro* and *in vivo* studies have strongly suggested that CD200 negatively regulates myeloid functions, particularly cells of the macrophage/dendritic cell lineage ^41^. The increased and specific expression of *Cd200* in IRCF makes tempting to hypothesize that its expression after MI may be implicated in controlling inflammation and immune response after MI ^34,42^.

Understanding the regulation of IRCF will be essential for identification of therapeutic strategies. Our transcriptomic studies together with our binding site analysis indicate a potential role for different transcription factors (*Sox9, Runx1* or *Smad*) in the generation of IRCF. In fact, these TFs have been previously implicated in cardiac fibrosis ^43,44^ and fibroblast regulation ^45^ although based on these studies and our own results it is unlikely that there is a single master regulator, but rather a combination of TFs (**Fig. 4b,c** **and Fig. S9a**). TGF-β signaling pathway represents the master regulator in activation of fibroblast and the development of cardiac fibrosis ^46,47^, where canonical signaling involves activation of SMAD2 and SMAD3 leading to promotion of myofibroblast formation and ECM production ^48–50^. Interestingly, our results demonstrate that the non-canonical TGF-β/PI3K-Akt signaling pathway also plays a significant role in the expression of *Cthrc1* and the activation of IRCF (**Fig. 4f** **and Fig. S9d**). Although further studies of the downstream mechanisms controlling these processes are required, our findings along with the development of specific inhibitors of the PI3K/Akt pathway could define new targets for modulating cardiac fibrosis after MI ^51^.

In conclusion, our study including the largest number of CF interrogated by scRNA-seq clearly demonstrates the remarkable heterogeneity of CF after MI and defines a specific subpopulation, characterized by the expression of specific markers. Insights provided into their regulation and the functional results indicating their role in the healing process after MI should help in the search for a more personalized treatment allowing the control of the fibrosis size in patients with MI.

## ACKNOWLEDGEMENTS

This work was supported by funds from the ISCIII and FEDER funds (PI16/00129, CPII15/00017), Red TERCEL RETIC RD16/0011/0005 and MINECO (Program RETOS Cardiomesh), ERANET II (Nanoreheart). AR-V is supported by FSE/Ministerio de Economía, Industria y Competitividad - Agencia Estatal de Investigación/ IJCI-2016-30254. N.F. is supported by a fellowship from the European Molecular Biology Organization (EMBO ALTF 241-2017). C.B. is supported by a New Frontiers Group award of the Austrian Academy of Sciences and by an ERC Starting Grant (EU Horizon 2020 research and innovation programme, grant agreement n° 679146). This work was supported in part, by the National Heart, Lung, and Blood Institute of the National Institute of Health under grant R01 HL136560

## AUTHOR CONTRIBUTIONS

**Conception and design**: AR-V, JPR, SCH, VL, DL-A, FP

**Development of methodology**: AR-V, JPR, SCH, AV-Z, PS-U, EL-V, BP

**Acquisition of data and assistance with experiments**: AR-V, SCH, MP, JJG, EI, GA, DA, GM, IP, YRJ, SR, HY, BP

**Analysis and interpretation of data**: AR-V, JPR, SCH, NF, LC, GB, XM-M, CB, DG-C, BP, VL, DL-A, FP

**Writing, review, and/or revision of the manuscript**: AR-V, JPR, SCH, NF, BP, VL, DL-A, FP

**Administrative, technical, or material support (i.e., reporting or organizing data, constructing databases)**: AV-Z, LC, PS-U, EL-V, PG-O, MP, JJG, GB, SJ, IP, YRJ, SR, HY

**Study supervision**: VL, DL-A, FP

## SUPPLEMENTARY FIGURE LEGENDS

**Supplementary Figure 1: Molecular and cellular characterization of *Col1α1-GFP* a)** Representative gating for isolation of GFP^+^/CD31^−^/CD45^−^ cardiac interstitial cells. *From left to right*: gating for singlets, viable cells (GFP^+^/TOPRO3^−^), metabolically active (Calcein^+^/TOPRO3^−^), and GFP^+^ cells that are negative for CD45 and CD31. **b)** Correlation heatmap of transcriptomic profiles between GFP^+^/CD31^−^/CD45^−^ cardiac interstitial cells, endothelial (CD31^+^), and bone marrow-derived (CD45^+^) cells, isolated from the heart (healthy, 7, 14 and 30 dpi), and dermal GFP^+^/CD31^−^/CD45^−^ fibroblasts from the tail (left). PCA scatter plot of cardiac interstitial, endothelial, bone marrow-derived and dermal samples subjected to transcriptome profiling (right). **c)** Boxplot of expression levels of traditional markers for non-activated (*Pdgfrα, Thy1, Ddr2, Tcf21, P4hb, Col1α1*) and activated CF (*Acta2, Postn, Ckap4, Sca1*) in different cell types.

**Supplementary Figure 2: Histological characterization of GFP^+^ cardiac interstitial cells from *Col1α1-GFP* a)** Percentage of GFP^+^/CD31^−^/CD45^−^ cells that express classical cell surface markers for CF (CD90 (THY1), mEFSK4, PDGFRα (CD140a)). **b)** Percentage of GFP^+^/CD31^−^/CD45^−^ cells in healthy myocardium and at 7, 14 and 30 dpi (left). Quantification of GFP^+^ cells at 7dpi (RZ, remote zone; BZ, border zone; IZ, infarct zone) and in healthy myocardium (RV, right ventricle; LV, left ventricle) (right). **c)** Immunofluorescence analysis of GFP (green), CD31 or CD45 (red), α-sarcomeric actin (grey) and DAPI/nuclei (blue) in the LV in healthy myocardium, and at 7, 14 and 30 dpi. Dotted lines delimit IZ from BZ. Epi, Epicardium; Endo, endocardium.

**Supplementary Figure 3: Quality control of single-cell RNA sequencing analysis a)** Scatter plot with number of UMIs (x-axis) and number of genes (y-axis) for the single cell analyses performed at different days after myocardial infarction. Color shows cells that passed quality filters (blue) and discarded ones (red). **b)** Histogram of the proportion of UMIs assigned to mitochondrial genes for the single cell analyses performed at different dpi. Color shows cells that passed quality filters (blue) and discarded ones (red). **c)** Boxplots of quality control variables (number of genes, number of UMIs and relative expression of mitochondrial genes) for the different single cell datasets.

**Supplementary Figure 4: Transcriptional signatures of the different clusters of GFP^+^ cardiac interstitial cells a)** Violin plots showing the normalized log expression (x-axis) of traditional markers for non-activated and activated CF (*Pdgfrα, Tcf21, Acta2, Thy1, Ckap4, P4hb, Ddr2, Sca1*) per cluster (A to K) from healthy, 7, 14 and 30 dpi (left). Violin plot showing the normalized log expression (x-axis) of the five top markers in cluster K for pericytes (*Rgs5, Higd1b, Vtn, Tinagl1, Colec11,*) per cluster (A to K) from healthy, 7, 14 and 30 dpi) (right). **b)** Violin plots of the normalized log expression of the top five markers per cluster. Color indicates the corresponding cluster (A to K). **c)** Boxplot denoting the specificity of distribution for the markers of every identified cluster. The specific (y-axis) value corresponds to the percentage of cells of a particular cluster expressing a given marker minus the percentage of every other cell expressing the same marker. **d)** Dot plot comparison of enriched GO terms for each of the identified clusters. The analysis was performed using the previously defined cluster markers.

**Supplementary Figure 5: Dynamics of the clusters during the cardiac healing process a)** Bar plots showing the percentage of cells (y-axis) per cluster relative to the total number of cells at each time point (grey scale). **b)** Dot plot comparison of expression and specificity of markers for non-activated fibroblasts (*Pdgfrα, Thy1, Ddr2, Tcf21, P4hb, Col1α1*), activated fibroblasts (*Acta2, Ckap4, Postn*) and cluster B markers (*Cthrc1, Comp, Ddah1, Sfrp2, Lox, Fmod*) in healthy heart, and at day 7, 14 and 30 dpi. Dot size represents percentage of cells, included in the cluster that express a given marker and the expression level defined by color intensity. **c)** Network representation of enriched pathways for the defined transcriptional waves (early, intermediate and late). Dot size represents the number of IRCFs markers annotated for each pathway and color the statistical significance.

**Supplementary Figure 6: Periostin (*Postn*) expression in CF at different time points after MI a)** t-SNE representation of the level of expression of *Postn* in healthy myocardium and at 7, 14 and 30 dpi (orange dots). **b)** Violin plots indicate level of expression of *Postn* in all CF in healthy myocardium and at 7, 14 and 30 dpi in all the clusters (above) and in cluster B (below). **c)** Staining of POSTN (red) in left (left) and right (right) healthy ventricles (above), and infarcted *vs* remote zones at 7, 14 and 30 dpi (GFP, green, arrowheads; DAPI/nuclei, blue) (below). Co-localization of GFP^+^ and POSTN^+^ in yellow (arrows). Epi, Epicardium.

**Supplementary Figure 7: Location of Cluster B specific CF at 7, 14 and 30 dpi** Overview of transversal sections of *Col1α1-GFP* infarcted hearts at 7, 14 and 30 dpi. Spatial location of CTHRC1, DDAH1, or FMOD in the infarct zone (IZ) and the remote zone (RZ). GFP^+^ (green, arrowheads), CTHRC1, DDAH1, or FMOD (red), Nuclei (DAPI, blue). Co-localizations are in yellow (arrows). Dotted lines delimits IZ. Epi, Epicardium; Endo, endocardium.

**Supplementary Figure 8: Expression of *Cd200* in Infarct Repair Cardiac Fibroblast a)** t-SNE projection (left) and violin plots (right) of normalized expression of *Cd146* (*Mcam*), *Cd200 (Ox2)* or *Cd146/Cd200* in all states. **b)** Representative gating for isolation of GFP^+^/CD200^+^/CD146^−^/CD31^−^/CD45^−^ and GFP^+^/CD200^−^/CD146^−^/CD31^−^/CD45^−^ cardiac interstitial cells. *From left to right*: gating for viable cells (7AAD^−^), singlets, GFP^+^ cells, CD45^−^ and CD31^−^, CD146^−^, and CD200^+^ or CD200^−^. **c)** Correlation of IRCF markers between single cell RNA-seq aggregated expression and bulk RNA-seq expression in CD200^+^ cells isolated at 7dpi.

**Supplementary Figure 9: Transcriptomic and functional regulation of IRCF a)** Violin plots representation of the expression of transcription factors (*Atf3, Bach2, Egr1, Fosb, Fosl1, Fosl2, Jun, Rela, Runx1, Rxra, Smad2, Smad3, Smad4, Sox9, Tcf3, Tead1, Tead2, Tead3, Tead4*) in all the clusters. **b)** Box plot representation of expression of IRCF markers after lentiviral infection with *Runx1* and *Sox9*. **c)** Western blot for p-Akt and total Akt protein cell lysates from CF cultured for 1 hour without treatment, DMSO, LY294002, TGF-β, TGF-β + DMSO and TGF-β + LY294002 **d)** Normalized expression bar plots for genes annotated to enriched GO terms (left) and top IRCF markers (right).

**Supplementary Figure 10: Characterization and transcriptomic analysis of *Cthrc1-KO* mice and CF a)** Normalized expression bar plots of expression levels for specific markers of cell types (*Col1α1, Col1α2, Myh6, Myh7 Cd31* (*Pecam1*)*, Cd45* (*Ptprc*)) (left) and for markers of cluster B (IRCF) (*Comp, Cthrc1, Ddah1, Fmod, Lox, Ptn*) (right) in different cell types (CD31, CD45, CF, Cardiomyocytes). **b)** Flow cytometry analysis of CTHRC1^+^ cardiac interstitial cells in WT and KO mice in healthy myocardium and at 3dpi (left). CD31 and CD45 expression in viable, CTHRC1^+^ cardiac interstitial cells in WT mice at 3 dpi (right). **c)** Western blot for Procollagen-I, CTHRC1 and β-Actin performed on cell lysates (CL) and in conditioned media (CM) from cultured E12.5 mouse embryonic fibroblasts (MEFs) of KO and WT mice. **d)** Localization of CTHRC1 (green, arrows) in the left ventricle of healthy WT mice, and in KO and WT mice at 3 and 5 dpi. Cardiac Troponin-I^+^ cardiomyocytes (red), Nuclei (DAPI, blue). Dotted lines delimits BZ. **e)** Spatial location of DDAH1, FMOD, CD31 and CD45 in the infarct zone (IZ) of WT and *Cthrc1-KO* mice at 5dpi. DDAH1 or FMOD (green, arrowheads), CD31 or CD45 (red, asterisks), Nuclei (DAPI, blue). Co-localizations are in yellow (arrows).

## REFERENCES

1. Pinto, A.R., et al. Revisiting Cardiac Cellular Composition. Circ Res 118, 400–409 (2016).

2. Frangogiannis, N.G. Pathophysiology of Myocardial Infarction. Compr Physiol 5, 1841–1875 (2015).

3. Shinde, A.V. & Frangogiannis, N.G. Mechanisms of Fibroblast Activation in the Remodeling Myocardium. Curr Pathobiol Rep 5, 145–152 (2017).

4. Tallquist, M.D. & Molkentin, J.D. Redefining the identity of cardiac fibroblasts. Nat Rev Cardiol 14, 484–491 (2017).

5. Zeisberg, E.M., et al. Endothelial-to-mesenchymal transition contributes to cardiac fibrosis. Nat Med 13, 952–961 (2007).

6. Mollmann, H., et al. Bone marrow-derived cells contribute to infarct remodelling. Cardiovasc Res 71, 661–671 (2006).

7. Moore-Morris, T., et al. Resident fibroblast lineages mediate pressure overload-induced cardiac fibrosis. J Clin Invest 124, 2921–2934 (2014).

8. Ruiz-Villalba, A., et al. Interacting resident epicardium-derived fibroblasts and recruited bone marrow cells form myocardial infarction scar. J Am Coll Cardiol 65, 2057–2066 (2015).

9. Kanisicak, O., et al. Genetic lineage tracing defines myofibroblast origin and function in the injured heart. Nature communications 7, 12260 (2016).

10. Frangogiannis, N.G. The role of transforming growth factor (TGF)-beta in the infarcted myocardium. J Thorac Dis 9, S52–S63 (2017).

11. Pyagay, P., et al. Collagen triple helix repeat containing 1, a novel secreted protein in injured and diseased arteries, inhibits collagen expression and promotes cell migration. Circ Res 96, 261–268 (2005).

12. Jin, Y.R., et al. Inhibition of osteoclast differentiation and collagen antibody-induced arthritis by CTHRC1. Bone 97, 153–167 (2017).

13. Yata, Y., et al. DNase I-hypersensitive sites enhance alpha1(I) collagen gene expression in hepatic stellate cells. Hepatology 37, 267–276 (2003).

14. Skelly, D.A., et al. Single-Cell Transcriptional Profiling Reveals Cellular Diversity and Intercommunication in the Mouse Heart. Cell Rep 22, 600–610 (2018).

15. Horn, M.A. & Trafford, A.W. Aging and the cardiac collagen matrix: Novel mediators of fibrotic remodelling. J Mol Cell Cardiol 93, 175–185 (2016).

16. Gonzalez, A., Schelbert, E.B., Diez, J. & Butler, J. Myocardial Interstitial Fibrosis in Heart Failure: Biological and Translational Perspectives. J Am Coll Cardiol 71, 1696–1706 (2018).

17. Kuhn, B., et al. Periostin induces proliferation of differentiated cardiomyocytes and promotes cardiac repair. Nat Med 13, 962–969 (2007).

18. Shimazaki, M., et al. Periostin is essential for cardiac healing after acute myocardial infarction. J Exp Med 205, 295–303 (2008).

19. Binks, A.P., Beyer, M., Miller, R. & LeClair, R.J. Cthrc1 lowers pulmonary collagen associated with bleomycin-induced fibrosis and protects lung function. Physiol Rep 5(2017).

20. LeClair, R.J., et al. Cthrc1 is a novel inhibitor of transforming growth factor-beta signaling and neointimal lesion formation. Circ Res 100, 826–833 (2007).

21. Stohn, J.P., Perreault, N.G., Wang, Q., Liaw, L. & Lindner, V. Cthrc1, a novel circulating hormone regulating metabolism. PLoS One 7, e47142 (2012).

22. Spinale, F.G. Matrix metalloproteinases: regulation and dysregulation in the failing heart. Circ Res 90, 520–530 (2002).

23. Huebener, P., et al. CD44 is critically involved in infarct healing by regulating the inflammatory and fibrotic response. J Immunol 180, 2625–2633 (2008).

24. Ivey, M.J. & Tallquist, M.D. Defining the Cardiac Fibroblast. Circ J 80, 2269–2276 (2016).

25. Borriello, F., Lederer, J., Scott, S. & Sharpe, A.H. MRC OX-2 defines a novel T cell costimulatory pathway. J Immunol 158, 4548–4554 (1997).

26. Hu, H.H., et al. New insights into TGF-beta/Smad signaling in tissue fibrosis. Chem Biol Interact 292, 76–83 (2018).

27. Montiel-Duarte, C., et al. Resistance to Imatinib Mesylate-induced apoptosis in acute lymphoblastic leukemia is associated with PTEN down-regulation due to promoter hypermethylation. Leuk Res 32, 709–716 (2008).

28. Kretzschmar, K., et al. Profiling proliferative cells and their progeny in damaged murine hearts. Proc Natl Acad Sci U S A 115, E12245–E12254 (2018).

29. Gladka, M.M., et al. Single-Cell Sequencing of the Healthy and Diseased Heart Reveals Cytoskeleton-Associated Protein 4 as a New Modulator of Fibroblasts Activation. Circulation 138, 166–180 (2018).

30. Farbehi, N., et al. Single-cell expression profiling reveals dynamic flux of cardiac stromal, vascular and immune cells in health and injury. Elife 8(2019).

31. Fu, X., et al. Specialized fibroblast differentiated states underlie scar formation in the infarcted mouse heart. J Clin Invest 128, 2127–2143 (2018).

32. Buch, T., et al. A Cre-inducible diphtheria toxin receptor mediates cell lineage ablation after toxin administration. Nat Methods 2, 419–426 (2005).

33. Li, J., et al. Collagen triple helix repeat containing-1 inhibits transforming growth factor-b1-induced collagen type I expression in keloid. Br J Dermatol 164, 1030–1036 (2011).

34. Prabhu, S.D. & Frangogiannis, N.G. The Biological Basis for Cardiac Repair After Myocardial Infarction: From Inflammation to Fibrosis. Circ Res 119, 91–112 (2016).

35. Kaur, H., et al. Targeted Ablation of Periostin-Expressing Activated Fibroblasts Prevents Adverse Cardiac Remodeling in Mice. Circ Res 118, 1906–1917 (2016).

36. Stohn, J.P., et al. Cthrc1 controls adipose tissue formation, body composition, and physical activity. Obesity (Silver Spring) 23, 1633–1642 (2015).

37. Duarte, C.W., et al. Elevated plasma levels of the pituitary hormone Cthrc1 in individuals with red hair but not in patients with solid tumors. PLoS One 9, e100449 (2014).

38. Wright, G.J., et al. Characterization of the CD200 receptor family in mice and humans and their interactions with CD200. J Immunol 171, 3034–3046 (2003).

39. Ishibashi, M., et al. CD200-positive cancer associated fibroblasts augment the sensitivity of Epidermal Growth Factor Receptor mutation-positive lung adenocarcinomas to EGFR Tyrosine kinase inhibitors. Sci Rep 7, 46662 (2017).

40. Liu, J.Q., et al. A Critical Role for CD200R Signaling in Limiting the Growth and Metastasis of CD200+ Melanoma. J Immunol 197, 1489–1497 (2016).

41. Jenmalm, M.C., Cherwinski, H., Bowman, E.P., Phillips, J.H. & Sedgwick, J.D. Regulation of myeloid cell function through the CD200 receptor. J Immunol 176, 191–199 (2006).

42. Nah, D.Y. & Rhee, M.Y. The inflammatory response and cardiac repair after myocardial infarction. Korean Circ J 39, 393–398 (2009).

43. Lacraz, G.P.A., et al. Tomo-Seq Identifies SOX9 as a Key Regulator of Cardiac Fibrosis During Ischemic Injury. Circulation 136, 1396–1409 (2017).

44. Bujak, M., et al. Essential role of Smad3 in infarct healing and in the pathogenesis of cardiac remodeling. Circulation 116, 2127–2138 (2007).

45. Kim, W., et al. RUNX1 is essential for mesenchymal stem cell proliferation and myofibroblast differentiation. Proc Natl Acad Sci U S A 111, 16389–16394 (2014).

46. Liu, G., Ma, C., Yang, H. & Zhang, P.Y. Transforming growth factor beta and its role in heart disease. Exp Ther Med 13, 2123–2128 (2017).

47. Dobaczewski, M., Chen, W. & Frangogiannis, N.G. Transforming growth factor (TGF)-beta signaling in cardiac remodeling. J Mol Cell Cardiol 51, 600–606 (2011).

48. Khalil, H., et al. Fibroblast-specific TGF-beta-Smad2/3 signaling underlies cardiac fibrosis. J Clin Invest 127, 3770–3783 (2017).

49. Frangogiannis, N.G., et al. Critical role of endogenous thrombospondin-1 in preventing expansion of healing myocardial infarcts. Circulation 111, 2935–2942 (2005).

50. Lal, H., et al. Cardiac fibroblast glycogen synthase kinase-3beta regulates ventricular remodeling and dysfunction in ischemic heart. Circulation 130, 419–430 (2014).

51. Campa, C.C., et al. Inhalation of the prodrug PI3K inhibitor CL27c improves lung function in asthma and fibrosis. Nature communications 9, 5232 (2018).

