## Supplementary material for "Single-cell RNA-seq analysis reveals the crucial role of Collagen Triplex Helix Repeat Containing 1 (CTHRC1) cardiac fibroblasts for ventricular remodeling after myocardial infarction": Material and methods

##### Animal models

All animal requisitions, housing, treatments and procedures have conformed to all state and institutional laws, guidelines and regulations and has been approved by Ethics Committee for Animal Research University of Navarra and the Government of Navarra. *Collagen1α1-GFP* and *Cthrc1-KO* mice were previously described [1,2].

*Induction of MI in mice*

Myocardial infarction (MI) was induced in mice by ligation of the left anterior descending (LAD) coronary artery as previously described [3]. Briefly, 8 to 10 weeks old mice were anesthetized through vaporized isoflurane, orally tubed using a 20G intravenous catheter, mechanically ventilated, and placed on a heating pad to maintain body temperature. A left thoracotomy was performed at the fourth-fifth intercostal space, where muscles were dissected. The LAD coronary artery was permanently ligated using a 7/0 non-absorbable ethylene suture. After visual verification of anemia and akinesis of the apex and anterior-lateral wall to ensure coronary occlusion, the thorax was closed in layers. After de-intubation, mice were kept warm until fully recovered.

*Cardiac function evaluation in mice*

Echocardiography was performed using a Vevo 2100 ultrasound system (VisualSonics, Toronto, Canada) with a 40-MHz transducer before ligation and at 7dpi. Measurements were optimized for small animals and performed as previously described [4]. Left ventricular M-mode tracings were obtained at the level of papillary muscles using the two-dimensional parasternal short axis imaging as a guide. Data was analyzed by the heart function package provided by VisualSonics Systems. Diastolic and systolic left ventricular (LV) internal dimensions (LVIDd and LVIDs) and fractional shortening (FS) (%) were measured to evaluate LV structural and functional changes. Statistical significance was analyzed by an unpaired Student's *t*-test using GraphPad Prim 6.0 (GraphPad Software) ($p<0.05$).

*Induction of MI in pig*

Experimental myocardial infarction was performed in approximately 12-15 month-old young adult male Göttingen minipigs weighing 20-30 kg (Ellegaard; Soroe Landevej 3024261, Dalmose, Denmark).

Pigs were premedicated during 14 days before the infarct with Amiodarone (400mg/day for the first week and 200 mg/day for the second week) and D-1 with Aspirin (300 mg) and Clopidrogel (300 mg). For the infarct procedure, animals were weighed and sedated with a mixture of tiletamine and zolepam (Zoletil®) 5 mg/kg i.m. Following adequate sedation, the neck was shaved, properly scrubbed and disinfected with Povidone Iodure and alcohol. Then, the pig was transferred to the surgery room and placed on the operating table. One intravenous catheter was placed in a marginal vein of one ear. An appropriately sized endotracheal tube was inserted for mechanical ventilation. Body temperature was maintained at 36.5-39 °C during the procedure with a blanket. Surface electrocardiogram (ECG) (lead II) was placed and connected to a DASH 5000 to monitor the onset of arrhythmias (ventricular tachycardia and fibrillation). To induce a stable and prolonged anesthesia, i.v. propofol infusion was initiated once the pig was under ventilation (30% $O_2$, ~400 mL tidal volume, ~18 breaths per min). Propofol dosing was adjusted to ~10 mg/kg per hour throughout the experimental procedure, depending on the anesthetic state of each animal. Adequate anesthesia was checked continuously. The animal was fitted with sterile drapes and sterile instruments were used by the surgeon. All surgical procedures were performed using standard sterile techniques, including sterile gowns, gloves, masks and instruments. Remifentanil 18-20 μg/kg/hr IV was given for intraoperative analgesia.

A small incision was performed in the neck in order to expose the carotid artery. Using the Seldinger technique, an introducer needle was placed into the carotid. Under fluoroscopy guidance, a JR 3.5 catheter was advanced over the wire to the level of the coronary sinus and place at the coronary ostium without full engagement. Good placement was confirmed with a bolus of contrast angiogram (VISIPAQUE) to visualize the left main, circumflex and anterior descending coronary. Heparin 300 UI/kg i.v. was given to prevent clotting just before LAD coronary artery occlusion. A guide wire was advanced into the LAD coronary artery and a balloon catheter (3.5 x 15-20) was advanced just below the first diagonal and inflated at 4-6 atm to ensure occlusion for 150 min of the LAD coronary artery confirmed by coronary angiogram. Then the balloon catheter was deflated, slowly removed and reperfusion documented by coronary angiography. In case of ventricular fibrillation, defibrillation was immediately performed using a HP defribillator. A bolus of lidocaine (2 mg/kg i.v.) and nitroglycerin (40 μg/kg intracoronary) was administrated to avoid vasospasms and arrhythmia. During the recovery period, phentanyl patches (0.03 mg/kg transdermal) was administrated for analgesia. Animals slowly woke-up under warming lamp in individual cage and extubation was performed when the pig begin to chew. Amoxicillin (Dufamox® 0.015 mg/kg i.m. antibiotic) and Carprofen (2 mg/kg s.c. NSAIDS) was given for post-procedural care.

At 6 months′ post-ischemia, animals were sacrificed with pentobarbital and a saturated solution of potassium chloride, hearts were excised and cut in 6 transversal sections (**Fig. 6a**). Half of the sections (even) were fixed in Zn-Formalin, paraffin embedded and sectioned for histological assessment. The other half (odd) were dissected according to the anatomical region of infarction (IZ, BZ, RZ) in four representative 1-2 $cm^2$ blocks and snap-frozen for following molecular assessments.

Pig samples from hearts with a subacute infarct were kindly provided by the laboratory of Professor S. Janssen (Univ. of Leuven). Adult domestic farm pigs (body weight: 30-34 kg) were used for the study. Briefly, under fluoroscopic guidance, a 7-French guiding catheter was delivered through the left coronary ostium just below the left anterior descendent distal to first diagonal branch and a balloon catheter was inflated, producing the MI. Occlusion was maintained for 75 min. After deflation, restoration of LAD blood flow was verified by coronary angiography and ST segment changes by ECG. Eight days after MI induction, animals were sacrificed and heart biopsies from IZ and RZ of the heart were collected for molecular and histological assessment.

Human heart samples were kindly provided by Professor S. Janssen. Heart biopsies from the right and left ventricle of non-infarcted (77y female) and chronically infarcted patients (55y, 69y males) were collected for molecular and histological analysis.

*Cardiac function evaluation in pig*

The week before induction of MI and 180dpi (end of study) animals were followed-up by echocardiography and magnetic resonance. Briefly, following adequate sedation with Zoletil® and induction of a stable and prolonged anesthesia, isofluorane inhalator, the left side of the chest was shaved and properly cleaned. Echocardiographic examinations were carried out using a cardiac ultrasound system equipped with a 4 MHz transducer (Philips SONOS 5500 M 2424A). The echocardiographic images were obtained using a right approach (4th or 5th intercostal space, in-left decubitus) for obtaining images of the parasternal long-axis and short-axis views. Three consecutive acquisitions for each view were performed and analyzed by two independent trained readers. Left ventricular volumes were calculated using the Simpson's method. Stroke volume was determined as: SV (mL) = LV diastolic volume – LV systolic volume). Ejection fraction was determined as EF (%) = (SV / LV diastolic volume) x 100 and cardiac output as CO (mL/min) = SV x heart rate.

Cardiac magnetic resonance examinations were performed on a 1.5 Tesla system (MAGNETOM Symphony with Tim, Siemens Healthcare GmbH, Erlangen, Germany) equipped with commercially available cardiac MRI software, electrocardiographic triggering and cardiac-dedicated surface coils. Pigs were placed in the left lateral decubitus position. Images were acquired with mechanical ventilation. MRI-compatible ECG pads were placed on the chest, using the guidelines of the vendor, and connected to the scanner. The ECG waveforms were examined for a clear R wave and adjusted if necessary. Standard steady-state free-precession (SSFP) cine images were acquired in the four-chamber, two-chamber and short-axis views to assess left-ventricular function with the following parameters: 8 mm slice thickness, no gap, TR= 43.26 ms, TE= 1.3 ms, flip angle= 80, 156 x 192 matrix, field of view of 260-280 x 325-375 mm, 1.7 x 1.7 mm pixel size, 14 segments, and 25 phases. To assess myocardial area at risk, a T2-STIR sequence was used in the short axis view. Rest perfusion was performed after the injection of 0.05 mmol/kg of intravenous contrast (gadobutrol, Gadovist, Bayer Schering Pharma AG, Berlin-Wedding, Germany) using an automated dual-head injector (Medrad Inc, Warrendale, Pennsylvania, USA) at 4 mL/s. Additional 0.1 mmol/kg of gadolinium was administered (total dose of 0.15 mmol/kg) before acquiring early and delayed gadolinium enhanced images, for which a turbo-flash phase-sensitive inversion recovery (PSIR) sequence (FA= 25°, TR/TE= 53.68/3.19 ms, voxel size= 1.4 × 1.4 × 8 mm) was employed. Sequence parameters may be modified if necessary.

**Single cell preparation and flow cytometry analysis**
Mouse cardiac interstitial cells (CIC) were obtained from 8-10 week old mice as previously described [3]. Briefly, after sacrifice the thorax was opened and the heart was perfused with ice-cold phosphate buffered saline pH 7.6 (PBS) (Lonza), atria were excised and, when suitable, hearts were divided up into remote, border and infarct zone (i.e. for zonal RNA-seq) **(Fig. 2F)**. Excised tissues were placed in DMEM medium (Sigma) supplemented with 10% FBS (Hyclone, GE) on ice. Next, ventricles were minced using a sterile scalpel. Pieces of tissue were incubated in Liberase TH (125 µg/mL) (Roche) in HBSS++ solution (Hanks balanced salt solution, Gibco) for 10 min at 37 °C in an orbital shaker. After the enzymatic incubation the partially digested tissue was mechanically dissociated by slowly pipetting to reach a single cell suspension. The supernatant was filtered through a cell strainer to discard cardiomyocytes (40 µm, nylon; Falcon). The digestion was repeated with the sedimented pieces and the supernatants were pooled together. Erythrocytes were removed using RBC lysis buffer (eBioscience). The total time for enzymatic digestion was 30 min.

For cellular characterization of *Col1α1-GFP*$^{+}$ CIC cells, the pellet was resuspended in 100 μL of sorting buffer (2 mM EDTA (Invitrogen), 0.5% BSA (Sigma) in PBS) before staining with the corresponding antibodies (**Supplementary Table 1**) and reagents for flow cytometry or fluorescence activated sorting. After 15 min incubation at room temperature in the dark, calcein-violet (Life Technologies) and/or Vybrant® DyeCycle™ Orange (VDO, Life Technologies) were added to antibody/cell suspensions at final concentrations of 1 and 1.25 μM, respectively. After incubation with dyes in a 37 °C water bath for 10 min, samples were washed twice with sorting buffer and spun at 1,200 rpm for 5 min each time. Then, supernatant was discarded and the final pellet was resuspended in 250 μL of sorting buffer. Just before flow cytometry TO-PRO-3 (Invitrogen) was added to each sample for viability check. Events were gated for metabolically active (calcein$^{+}$), nucleated (VDO$^{+}$), and viable (TO-PRO-3$^{-}$) single cells **(Fig. S1a)**, before gating on CD31$^{-}$/CD45$^{-}$ events for the analysis of GFP$^{+}$ cardiac fibroblasts (CF), CD31$^{+}$/CD45$^{-}$ for endothelial cells, and CD45$^{+}$/CD31$^{-}$ for bone marrow-derived cells **(Fig. S1a)**. Flow cytometry for these samples was conducted on a FACSCANTO Flow Cytometer (BD Biosciences). The files generated by flow cytometry were processed using FlowJo Software (Tree Star, Ashland, USA). Statistical significance was analyzed by the Student's *t*-test, and shown as mean plus standard deviation using GraphPad Prim 6.0 (GraphPad Software) ($p<0.05$).

For cellular characterization of *Cthrc1-KO* and *Cthrc1-WT* CIC cells, pellet was resuspended in 100 μL of sorting buffer and treated with Human TruStain fcX™ (anti-mouse 16/32 antibody, BioLegend, San Diego, CA) to prevent non-specific binding, followed by incubation with relevant antibodies for 25 min at 40 °C. Cell-surface antigen expression was examined for CD31$^{+}$ and CD45$^{+}$ cells. For intracellular CTHRC1 staining, cell suspension was incubated in DMEM containing 5% FBS and 3 μg/mL Brefeldin A (BioLegend, San Diego, CA) for 5 hours. After treatment with a Cytofix/Cytoperm kit (BD Biosciences), the permeabilized cells were stained with polyclonal anti-CTHRC1 antibody in combination with PE-conjugated anti-rabbit IgG (Jackson ImmunoResearch Laboratories, Inc). Viable and non-viable cells were distinguished using LIVE/DEAD® Fixable Violet Stain kit (Life Technologies, Carlsbad, CA). Flow cytometry for these samples was conducted on a MacsQuant Analyzer 10 (Miltenyi Biotec., Inc.) and the data was processed using WinList 5.0 software (Verity software House).

**Cardiomyocytes isolation**

Five days after infarct induction, cardiomyocytes were isolated from hearts of infarcted as well as healthy mice. Animals were anesthetized, their chest was open and the heart cannulated for standard Langendorff retrograde perfusion [5]. Perfusion was performed under constant pressure (60 mmHg; 37 °C, 8 min) in $Ca^{2+}$-free buffer containing 113 mM NaCl, 4.7 mM KCl, 1.2 mM $MgSO_4$, 5.5 mM glucose, 0.6 mM $KH_2PO_4$, 0.6 mM $Na_2HPO_4$, 12 mM $NaHCO_3$, 10 mM $KHCO_3$, 10 mM Hepes, 10 mM 2,3-butanedione monoxime, and 30 mM taurine. Digestion was initiated by adding a mixture of recombinant enzymes (0.2 mg/mL Liberase Blendzyme (Roche), 0.14 mg/mL trypsin (Invitrogen)), and 12.5 μM $CaCl_2$ to the perfusion solution. When the heart became swollen (10 min), it was removed and gently teased into small pieces with fine forceps in the same enzyme solution. Heart tissue was further mechanically dissociated using 2, 1.5, and 1 mm-diameter pipettes, until all large heart tissue pieces were dispersed. The digestion buffer was neutralized with stopping buffer containing 10% FBS and 12.5 μM

$CaCl_2$. Cardiomyocytes were pelleted by gravity (20 min), the supernatant aspirated and cells resuspended in the perfusion solution containing 5% FBS and 12.5 μM $CaCl_2$. The calcium concentration was increased by gradually adding $CaCl_2$ from 62 μM to 1 mM final concentration. Cells were collected by centrifugation and the pellet was homogenized with TRIzol™ Reagent (ThermoFisher). RNA isolation was performed according to the manufacturer's instructions.

**Cell sorting**

CIC pellets from each heart were resuspended in 80 μL sorting buffer and incubated with 20 μL of Feeder Removal MicroBeads (Miltenyi) for 15 min at 4 °C. A positive selection of CIC, enriched in CF, was performed twice using LS columns (Miltenyi) according to the manufacturer's instructions. The negative fraction was analyzed to confirm that no $GFP^+$ cells were present. The positive fraction was centrifuged at 1,200 rpm and the pellet was resuspended in 100 μL of sorting buffer. Cells were incubated with the corresponding antibodies for 15 min at room temperature in the dark **(Supplementary Table 1)**. After incubation, the samples were washed twice with sorting buffer and spun at 1,200 rpm for 5 min each time, supernatant was discarded and the final pellet was resuspended in 250 μL sorting buffer. The cell sorting was performed using a FACSAria (BD Biosciences) and analyzed with FACSdiva software (BD Biosciences). Standard, strict forward scatter height *versus* area criteria were used to discriminate doublets and gate only singleton cells. Viable cells were identified as positive by staining with 7-AAD (BD Bioscience). Viable cells gated on the FSC/SCC were sorted on the basis of the expression of GFP, and/or staining with anti-feeder cells antibody (mEFSK4 clone) (Miltenyi).

**Single-cell RNA-sequencing (scRNA-seq)**

The transcriptome of isolated *$GFP^+$-CFs* from 8-12 weeks old, healthy mice or infarct mice at 7, 14 or 30 dpi were examined using Single Cell 3' Reagent Kits v2 (10X Genomics) according to the manufacturer's instructions. Two hearts were pooled in each scRNA-seq experiment. For *Cthrc1-KO* mice experiment only one heart was used. Briefly, 25,000 $GFP^+$/$CD31^-$/$CD45^-$ events were sorted in 1X PBS, 0.05% BSA and the number of cells was quantified in a Neubauer chamber. Approximately 16,000 cells were loaded at a concentration of 1,000 cells/μL on a Chromium Controller instrument (10X Genomics) to generate single-cell gel bead-in-emulsions (GEMs). In this step, each cell was encapsulated with primers containing a fixed Illumina Read 1 sequence, followed by a cell-identifying 16 bp 10X barcode, a 10 bp Unique Molecular Identifier (UMI) and a poly-dT sequence. A subsequent reverse transcription yielded full-length, barcoded cDNA. This cDNA was then released from the GEMs, PCR-amplified and purified with magnetic beads (SPRIselect, Beckman Coulter). Enzymatic Fragmentation and Size Selection was used to optimize cDNA size prior to library construction. Illumina adaptor sequences were added and the resulting library was amplified via end repair, A-tailing, adaptor ligation and PCR. Library quality control and quantification was performed using Qubit 3.0 Fluorometer (Life Technologies) and Agilent's 4200 TapeStation System, respectively. Sequencing was performed in a NextSeq500 (Illumina) (Read1: 26 cycles; Read2: 57 cycles; i7 index: 8 cycles) at an average depth of 50,000 reads/cell.

**Bulk RNA-sequencing (RNA-seq)**

Bulk RNA-seq was performed following MARS-seq protocol adapted for bulk RNA-seq [6,7] with minor modifications. Briefly, 3,000 to 10,000 cells were sorted in 100 μL of

Lysis/Binding Buffer (Ambion), vortexed and stored at -80 °C until further processing. Poly-A RNA was extracted with Dynabeads Oligo (dT) (Ambion) and reverse-transcribed with AffinityScript Multiple Temperature Reverse Transcriptase (RT) (Agilent) using poly-dT oligos (IDT) carrying a 7 bp index. Up to 8 samples with similar overall RNA content were pooled together and subjected to linear amplification via IVT using HiScribe T7 High Yield RNA Synthesis Kit (New England Biolabs (NEB)). The resulting antisense RNA was fragmented into 250-350 bps fragments with RNA Fragmentation Reagents (Ambion) and dephosphorylated with 1U FastAP (Thermo Scientific) for 15 min at 37 °C. Partial Illumina adaptor sequences [6] were ligated with T4 RNA Ligase 1 (NEB) followed by a second reverse transcription. Full Illumina adaptor sequences were added during library amplification with KAPA HiFi DNA Polymerase (Kapa Biosystems). Libraries were quantified using a Qubit 3.0 Fluorometer and their size profiles examined in Agilent's 4200 TapeStation System. Sequencing was carried out in an Illumina NextSeq 500 using paired-end, dual-index sequencing (Rd1: 68 cycles; Rd2: 15 cycles; i7: 8 cycles) at a depth of 10 million reads per sample.

**mcSCRB-seq**
The transcriptome of mouse cardiomyocytes was examined using mcSCRBseq [8], adapted for bulk RNA-seq (this paper). A 100 ng aliquot of total RNA was used in each experiment. Poly-A RNA was extracted with Dynabeads Oligo (dT) (Ambion) and eluted in 8 µL Tris Cl pH 7.5. Prior to denaturing at 72 °C for 3 min, 1 µL of 2 µM barcoded oligo-dT primer was added [8]. Reverse transcription was performed at 42 °C for 90 min in a mixture containing: 1X Maxima RT buffer, 7.5 % PEG 8000, 1 mM dNTPs, 5 µM of unblocked template-switching oligo [8], and 240 U of Maxima H, Minus Reverse Transcriptase (Thermo Scientific). Primer excess was removed through digestion with Exonuclease I (NEB). cDNA was purified with a 1.2X SPRI clean up (Agencourt AMPure XP, Beckman Coulter), then subjected to PCR amplification using Terra polymerase (Takara Bio) and SINGV6 primer with the following thermal cycling: 3 min at 98 °C followed by 6 cycles of 15 sec at 98 °C, 30 sec at 65 °C, and 4 min at 68 °C, and a final elongation step of 10 min at 72 °C. Upon purification (1X SPRI clean-up), 0.8 ng of double stranded cDNA were tagmented using Nextera-XT Tn5 (Illumina). PCR amplification using P5NEXTPT5 primer and i7 indexed primers allowed 3' enrichment and secondary barcoding of the libraries as well as addition of Illumina adaptor sequences. Final libraries were quantified and their profiles examined as described above. Sequencing was carried out in an Illumina NextSeq 500 using paired-end, dual-index sequencing (Rd1: 16 cycles; Rd2: 67 cycles; i7: 8 cycles) at a minimum depth of 10 million reads per sample.

**Truseq RNA-sequencing**
Total RNA from frozen biopsies of the different anatomical regions coming from swine and human samples was isolated using TRIzol reagent (Ambion). Following mechanical homogenization with an Ultra-turrax (T10 basis Ultra-Turrax, IKA), RNA was either extracted according to manufacturer's instructions or stored at -80 °C until processed. RNA concentration was quantified using a Qubit 3.0 Fluorometer and its quality was examined in Agilent's 4200 TapeStation System.

Swine cDNA libraries were performed using Truseq Stranded mRNA library prep kit (Illumina). mRNA was selected using poly-dT magnetic beads and followed by RT, second strand synthesis, 3' adenylation, Y shaped adaptor ligation and library enrichment. Custom Y shaped adaptors were used [9]. Final libraries were quantified and their profiles examined as described above. Sequencing was performed in an Illumina NextSeq 500 using single-end, dual-index sequencing (Rd1: 75 cycles; i7: 8 cycles; i5: 8 cycles) at a minimum depth of 10 million reads per samples.

Human RNA was subjected to rRNA depletion using Truseq Stranded Total RNA library prep Gold kit (Illumina), followed as described above, by RT, second strand synthesis, 3' adenylation, Y shaped adaptor ligation and library enrichment. Final libraries were quantified and their profiles examined as described above. Sequencing was performed in an Illumina NextSeq 500 using single-end, dual-index sequencing (Rd1: 75 cycles; i7: 8 cycles; i5: 8 cycles) at a minimum depth of 20 million reads per samples.

**ATAC-sequencing**
Accessible chromatin mapping was performed using FAST-ATAC-seq [10] with minor modifications. Briefly, 10,000 cells were sorted in 1X PBS, 0.05% BSA and pelleted by centrifugation at 500 rcf for 5 min at 4 °C with low acceleration and brake settings in a pre-cooled swinging-bucket rotor centrifuge. All the supernatant but 5 µL was removed. Then, 25 µL of transposase mix (15 µL of 2x TD buffer (Illumina), 1 µL of TDE1 (Illumina), 0.25 µL of 5% digitonin (Promega), 8.75 µL of nuclease-free water) were added to the cells and the pellet was disrupted by pipetting. Transposition reactions were incubated at 37 °C for 30 min in an Eppendorf ThermoMixer with shaking at 450 rpm. Reactions were stopped at 4 °C for 5 min. Next, in order to release tagmented DNA, samples were incubated at 40 °C for 30 min with 5 µL of clean up buffer (900 mM NaCl (Sigma), 30 mM EDTA (Millipore)), 2 µL of 5% SDS (Millipore) and 2 µL of Proteinase K (NEB). DNA was purified using a 2X SPRI beads cleanup (Agencourt AMPure XP, Beckman Coulter). In order to determine the total number of PCR cycles needed for library amplification, two sequential PCRs were performed using KAPA HiFi DNA Polymerase and customized Nextera PCR indexing primers (IDT) [11]. The conditions of the first PCR were: 5 min at 72 °C and 2 min at 98 °C followed by 9 cycles of 20 secs at 9 °C, 30 secs at 63 °C, and 1 min at 72 °C. Depending on the library concentration obtained in this first PCR, a second PCR (2 min at 98 °C followed by 4 to 6 cycles of 20 secs at 98 °C, 30 secs at 63 °C, and 1 min at 72 °C) was performed aiming for a library concentration in the range of 2 to 10 ng/µL. PCR products were purified using a 2X SPRI beads cleanup. Libraries were quantified and their size profiles examined as described above. Sequencing was carried out in an Illumina NextSeq 500 using paired-end, dual-index sequencing (Rd1: 38 cycles; Rd2: 38 cycles; i7: 8 cycles; i5: 8 cycles) at a depth of 80 million reads per sample.

**TGF-β signaling pathways *in vitro* functional assay**
For cell culture, CIC were resuspended in complete medium (DMEM (Sigma) + 10% FBS + 1% P/S (Life Technology) + 1% L-Glu (Life Technology)) containing 10 ng/mL of basic Fibroblasts Growth Factor (bFGF) (Peprotech) and plated in a 0.1% (w/v) gelatin-coated (Sigma) 6-well-plate well (one heart per well). CF were selected by attachment through washing the wells twice with PBS after overnight incubation and followed by a replacement of the complete medium plus 10 ng/mL of bFGF, lowering the concentration to 5 ng/mL (passage 1) and 2 ng/mL (passage 2) over time. After 24 hours in passage 2, cultured CF were incubated in starving conditions (complete DMEM without FBS) overnight, and then the medium was supplemented with hTGF-β1 (10 ng/mL, Peprotech), and LY294002 (40 µM, LC Laboratories) or DMSO (Sigma)

(used as vehicle) for 24-48 hours. To validate the effect of the inhibitor, LY294002, cell pellets were collected one hour after the treatments described above (TGF-β, TGF-β + DMSO, TGF-β + LY294002). Treated CF were submitted to bulk RNA-seq.

##### *In vitro* Scratch Assay

$GFP^+$ adult CF from three mice in passage 2 were seeded in 24-well culture plates at a density of 1 x $10^4$ cells/well. After 48 hours, cells were subjected to starving conditions overnight. Next day, a straight line was generated on the monolayer with a pipette tip. Cells were washed twice with PBS to remove debris and serum-free medium supplemented with different treatment (control, TGF-β, TGF-β + DMSO, TGF-β + LY294002) was added to the cells. The capacity of the cells to migrate and occupy the denuded area was tracked using time-lapse microscopy *Cell Observer Z1* (Zeiss) for 23 hours. Covered area was quantified using Fiji Software (1.46). Statistical significance was analyzed using a non-parametric one-way analysis of variance with a Kruskal-Wallis post-hoc test, and shown as mean plus standard deviation using GraphPad Prim 6.0 (GraphPad Software) ($p<0.05$).

##### Lentiviral vectors construction

Lentiviral vector containing *Runx1* cDNA (pLenti-GIII-CMV-Runx1-GFP-2A-puro) was purchased from ABM (LV496973). Cloning vector containing *Sox9* cDNA (pcDNA3.1+/C-(K)DYK, OMu22479D) was purchased from GenSpript for posterior cloning in the lentiviral vector. Briefly, *Sox9* cDNA was amplified from the pcDNA with primers containing restriction sites for the enzymes present in the pLenti around *Runx1*, *NheI* and *XbaI,* using Platinum®Taq DNA polymerase High Fidelity (Life Technologies), and cloned downstream of the CMV promoter in pLenti-GII-GFP-2A-Puro after restriction digestion of the vector and the amplification fragment with *NheI* and *XbaI* to yield pLenti-CMV-Sox9-GFP. Briefly, digestion was performed for 4 hours at 37 °C and fragments were either purified with NucleoSpin Gel and PCR Clean-up column (Macherey Nagel) directly or run in an electrophoresis gel and purified from the gel band with the same kit. Then, a ligation reaction was performed with the digested plasmid, the insert and 0.5 µL of T4 DNA ligase (NEB) and incubated overnight at 16 °C. Ligation product was transformed into 25 µL of Stbl3 competent cells (Life Technologies). Clones were tested by *NheI* and *XbaI* digestion and Sanger sequencing. An Empty lentiviral vector (pLenti-CMV-GFP) was constructed from the pLenti-GIII-CMV-Runx1-GFP-2A-puro, removing *Runx1* cDNA digesting the vector with *EcoRV* (NEB), present at both sides of the insert, purified and cloned as described above. Once cloned, constructs were confirmed by sequence.

##### Lentiviral production

All lentiviruses were produced for the infection of CFs by co-transfecting 5 x $10^6$ HEK-293T cells with 9 µg of transcription factor of interest encoding plasmid (*Runx1, Sox9, Empty*), 6 µg psPAX2 (Addgene, #12260) and 3 µg of pMD2G (Addgene, #12259) plasmids using 40 µL of Lipofectamine2000 (Invitrogen) and 1970 µL Opti-MEM (Gibco) in a 15 cm culture dish. Lentivirus-containing supernatant was harvested 72 hours later, filtered through a 0.45 µm HV-Durapore Stericups (SLHV033RS Millipore) and viral particles were purified and concentrated. Briefly, the filtrates were transferred to Ultra-Clear Beckman tubes (Beckman Coulters) and centrifuged at 26,000 rpm for 2.5 hours at 4 °C in an Ultracentrifuge (Beckman optima LE-80K). The pellet was resuspended in an appropriate volume of PBS depending on the number of culture dishes. The lentivirus stocks were snap-frozen in 100 µL aliquots and stored at -80 °C.

Vector titers were determined by serial dilutions of the concentrated lentivirus on HEK-293T cells in cultured medium supplemented with 8 μg/mL polybrene (Sigma) and analyzed 72 hours after infection by flow cytometry.

**Lentiviral transduction**

For transduction, CFs were infected at passage 1 with virus stock, according to the titer, in cultured medium supplemented with 8 μg/mL polybrene (Sigma). After addition of the virus, cell plates were centrifuged at 2,000 rpm for 90 min at 32 °C without acceleration or break in a pre-warmed swing rotor centrifuge. After 24 hours the infection was repeated. The cells were incubated for 72 hours from the first infection, they were subculture from passage 1 to 2. After four days in culture (passage 2) cells were trypsinized and sorted in 100 μL of Lysis/Binding Buffer, vortexed and stored at -80 °C until further processing. Cells were collected for RNA-seq and empty vector was used as control.

**Immunofluorescence**

Mice adult hearts were excised and washed in PBS, fixed in 4% fresh paraformaldehyde (Sigma), cryoprotected in sucrose (Sigma), embedded in OCT compound (VWR) and frozen in dry ice. Pig's biopsies of infarct, border and remote zones were paraffin embedded, sectioned and rehydrated. Five-to-ten micron sections were washed in Tris-PBS (TPBS), non-specific IgG binding sites blocked with 16% goat serum (Dako), 1% BSA, and 0.1% Triton X-100 (Sigma) (SBT) and incubated overnight in the corresponding primary antibodies at 4 °C (**Supplementary Table 3**). CTHRC1 and anti-GFP staining required an antigen retrieval step using 100 mM citrate buffer (Sodium citrate buffer and citric acid (Sigma)) before blocking. The sections were then washed and incubated with the appropriate fluorescence-conjugated secondary antibodies for 1 hour before mounting **(Supplementary Table 4)**. Negative controls were performed by incubating without the primary antibody. Cell nuclei were counterstained with 4',6-diamidino-2-phenylindole (DAPI) (Sigma) or Hoechst 33258 (Invitrogen) (1 μg/mL for 5 min). All images were captured in a Zeiss LSM 510 800 (Zeiss) or Leica SP8 confocal laser microcopy (Leica).

**Morphometric assay: quantification of GFP$^{+}$ and CD200$^{+}$/GFP$^{+}$ cells**

The quantification of the area occupied by GFP$^{+}$ or CD200$^{+}$/GFP$^{+}$ was performed in stained sections from healthy myocardium and infarct tissue at 7dpi. The number of pixels from GFP$^{+}$ (green) or CD200$^{+}$/GFP$^{+}$ (yellow) areas and DAPI$^{+}$ nuclei (blue) were determined using user manual macros in Fiji software (1.46). For infarcted hearts, we selected sections from infarct, border and remote zones (IZ, BZ, RZ, respectively). For healthy hearts, the free wall of the left ventricle (LV) and right ventricle (RV) were selected. Among three and four mice were used for the analysis. A total of three to four histological sections, collected every 150-200μm, were chosen for each anatomical region per animal. For graphic representation, the percentage of the area of GFP$^{+}$ or CD200$^{+}$/GFP$^{+}$ cells were calculated for each region. The results were analyzed using a non-parametric one-way analysis of variance with a Kruskal-Wallis post-hoc test using GraphPad Prism 6.0 (GraphPad Software) ($p<0.05$).

**Kaplan-Meier survival curve**

To analyze animal survival, MI was induced in groups of *Cthrc1-KO* and *Cthrc1-WT* animals (n=10 per group). Survival of animals was carefully monitored once a day for seven days after MI. Kaplan-Meier analyses and the log-rank Mantel-Cox test were used to determine statistical difference between the survival curves of the two groups of animals.

**Quantification of Collagen deposition**

Picrosirius red (PSR)-polarization detection of collagen fibers in tissue sections was used to determine collagen deposition in left ventricular walls at 3dpi *Cthrc1-KO* and *Cthrc1-WT* mice hearts (n= 5 per group). Slides were incubated with Sirius Red (0.1%) dissolved in aqueous saturated picric acid overnight. PSR-stained sections were imaged under polarized light using an Axioskop 40 microscope (Zeiss). Deposition of collagen fibers were analyzed using Fiji software (1.46). Statistical significance was analyzed by an unpaired Student's *t*-test using Graphpad Prism 6.0 (GraphPad Software) ($p<0.05$).

**Mouse embryonic fibroblasts culture**

Mouse embryonic fibroblast (MEFs) cultures were established from *Cthrc1-KO* and *Cthrc1-WT* 13.5 day embryos on the C57BL/6J background as described previously [12]. Cells were used between passages 2 to 5. Cell lysates and conditioned media were harvested for Western blotting 3 days after reaching confluence **(Fig. S10c)**.

**Western Blot**

Passage 2 CF, or MEFs were trypsinized, spun at 1,200 rpm 5 min, and pellets were frozen in liquid nitrogen and stored at -80 °C until further processing. Total protein was extracted using 1% of Triton-X100 within a cocktail of protease inhibitors (Complete-EDTA, Sodium orthovanadate, and Sodium fluoride (Sigma-Aldrich)), and quantified using BCA Protein Assay Kit (ThermoFisher Scientific). Proteins were separated using a 15% acrylamide gel and blotted in a nitrocellulose membrane (Bio-Rad, 0.45 µm). Membranes were blocked using a Blocking Solution (5% non-fat dry milk powder and 0.05% Tween-20 in TBS buffer (1 mM Tris-HCl, 15 mM NaCl)) for 1 hour at room temperature and incubated with the corresponding primary antibody (**Supplementary Table 2**) in Blocking Solution overnight at 4 °C. The membranes were washed using TBS-0.05% Tween-20 solution. Membranes were stripped using Restored™ Western Blot Stripping Buffer (0.5M, 10 min) (ThermoFisher Scientific), and incubated during 2 hours at room temperature with anti-β-actin or anti-α-tubulin (**Supplementary Table 2**) as internal reference. Lumigen ECL Ultra (TMA-6) (Beckman Coulter) was used for the development of the signal after incubation with the corresponding HRP-conjugated secondary antibody (**Supplementary Table 2**) for 5 min at room temperature. *ChemiDoc™ Imaging Systems* (Bio-Rad) imaging system was used for image generation.

**RNA isolation and qPCR**

Total RNA from frozen biopsies of the different anatomical regions was isolated as described above, using TRIzol reagent (Ambion) according to manufacturer's instructions. RNA purity (A260/A280 ratio $\geq$ 1.8) and concentration were measured using the Nanodrop (Thermo Fisher). Total RNA was stored at -80 °C. Prior to cDNA preparation, genomic DNA was removed using the Thermo Scientific DNase treatment kit (Thermo Fisher). A total of 1 µg of RNA was converted into cDNA using 125 µM of anchored oligo-dT primers (IDT) and Superscript II reverse transcriptase (RT) (Life Technologies). For each qPCR the cDNA equivalent to 5 ng RNA was used. The qPCR reactions contained power SYBR green PCR master mix (Applied Biosystems) and an equimolar primer mix (0.8 µM). The amplification protocol consisted of 2 min at 50 °C,

10 min at 95 °C, followed by 40 cycles of 15 secs at 95 °C, and 1 min at 60 °C, and completed with a standard melting curve protocol (15 secs at 95 °C, 1 min at 60 °C and 15 secs at 95 °C). The melting curve analysis (ViiA TM 7 Real-Time PCR system, Applied Biosystems) and size fractionation by agarose gel electrophoresis were used to confirm amplification of the expected products [13]. Target quantity ($N_0$) was obtained from the extracted raw data using *LinRegPCR* program [14]. Systematic differences induced by RT reactions in the observed expression levels per sample within tissue types as well as systematic differences between qPCR runs were removed using *Factor-qPCR* program [15]. *HPRT1* and *RPL4* were selected as reference genes for porcine samples after a stability analysis as described previously [16,17] using geNorm [18]. Primer sequences were designed using *primer3*, *BLAST* (NIH) and *oligo analyzer* (IDT) software, and are summarized in **Supplementary Table 5**. Graphs and statistical analysis were performed by a non-parametric one-way analysis of variance with a Kruskal-Wallis post-hoc test in *GraphPad* Prim version 6.0 (GraphPad Software) ($P<0.05$).

**scRNA-seq analysis**

Sequenced libraries were demultiplexed, aligned to the mouse transcriptome (*mm10*) and quantified using *Cell Ranger* (2.0.0) from 10X Genomics. The output of the pre-processing pipeline consisted of gene expression matrices per cell. Further computational analysis was performed using Seurat (v. 2.3.4). Cells were subjected to quality control filters based on the number of detected genes, number of UMIs and proportion of UMIs mapped to mitochondrial genes per cell. The thresholds for each of the single cell libraries were selected based on the distribution of the previously mentioned variables and visual inspection of quality control scatter plots **(Figure S3)**. Using these parameters, 7,079 (healthy myocardium), 10,448 (7dpi), 8,337 (14dpi), 6,805 (30dpi) and 4,189 (*Cthrc1-KO*) cells were retained.

Each single cell dataset was subjected to normalization, identification of highly variable genes and removal of unwanted sources of variation as described previously [19]. To integrate each of the analyzed time points, an analysis based on canonical correlation analysis (CCA) [20] was used. Briefly, genes identified among the top 1,000 most variable genes in at least 3 datasets were selected as input for CCA, and 30 canonical vectors were calculated. Sixteen CCA components were selected by visual inspection of biweight correlation plots. Cells whose variance explained by CCA was < 2-fold, compared with principal component analysis (PCA), were removed. The integrated single cell dataset was subjected to unsupervised clustering, with a resolution parameter of 0.8. Non-linear dimensional reduction was performed using t-distributed stochastic neighbor embedding (t-SNE) [21]. All differential expression tests were done using the function *FindAllMarkers* with default settings. Up-regulated genes detected in a fraction of 0.2 in either of the populations were deemed significant (p.value < 0.01).

**Bulk RNA-seq analysis**

Samples were demultiplexed using Illumina bcl2fastq software (1.2.4) and aligned to the mouse (*mm10*), swine (*Sscrofa 11.1*) and Human (*GRCh38*) genome with STAR (2.6.1) [22] setting the parameters to default values. Quantification and generation of gene expression matrices was performed with the function *featureCounts,* implemented in the R package Rsubread [23]. The *ensembl* transcriptomes (*GRCm38.91, Sscrofa11.1.93, and GRCh38.92*) were used as reference for gene annotation. Before statistical analysis, the function *filterbyExpr*, implemented in the R package *edgeR* [24], was used to determine genes with enough counts for further analyses. Data transformation, normalization and testing for differential expression was performed with DESeq2 [25].

**ATAC-seq analysis**

ATAC-seq libraries were demultiplexed using Illumina bcl2fastq software. The generated fastq files were aligned to the mouse (mm10) genome using bowtie2 (2.3.1) [26] with the parameters *--no-discordant, --no-mixed, --very-sensitive* and *-X 1000*. The generated SAM files were sorted and converted to BAM files using samtools (1.6). Peak calling was done using MACS2 (2.1.0) [27] with the parameters *-q 0.5, -B, --call-summit* and *--keep-dup=all*. Detected peaks were annotated to the *UCSC mm10* known gene database (3.4.4) using *ChIPpeakAnno* [28] R package. Peak quantification was done using *featureCounts* implemented in Rsubread [23]. Data transformation, normalization and testing for differential open chromatin regions was performed with DESeq2 [25]. We extracted differentially accessible regions between $CD200^{+}$ and $CD200^{-}$ by selecting the ones with Adjusted P.value < $1e^{-3}$ and LogFC > 0. Only distal regions (|Distance to TSS| > 3kb) were kept for further analyses. The selected regions were subjected to Motif Enrichment Analysis using *HOMER* (4.9). Novel and known motifs were identified using the function *findMotifsGenome.pl* with the parameter *-size 200*. All other parameters were set to default values

**Pathway and gene ontology analysis**

To perform enrichment analysis for signaling pathways and gene ontology categories, the R package *clusterProfiler* [29] was used. For pathway enrichment, we used the Reactome [30], and for gene ontology categories, we centered on the biological process ontology. Hypergeometric tests were performed and p-values were corrected using the Benjamini-Hochberg method [31], setting the p-value cutoff at 0.01 and the q-value cutoff at 0.05. Rest of the parameters were set to default values. Furthermore, we used enrichR as an interface to the *Enrichr* [32] database to perform additional tests for enrichment. In this case, the category *GO_Biological_Process_2018* was selected. Results were sorted according to the enrichr combined score, implemented in the *EnrichR* R package.

**Network betweenness analysis.**

To identify upstream regulators of gene expression changes mediated by TGF-β receptors, we generated a network spanning protein interactions and transcription factor-gene interactions. We downloaded transcription factors-genes interactions from *ENCODE*, *CHEA*, *ESCAPE*, *MotifMap*, *Transfac* using the *Harmonizome* [33] database (July 3, 2017). In addition, we downloaded the TTRUST [34] database (July 3, 2017). Gene names were mapped using NCBI (ftp://ftp.ncbi.nlm.nih.gov/gene/DATA/gene_info.gz (downloaded on October 10, 2016). Then the transcription factors with the strongest evidence were connected to each gene. 745 highly biased genes were connected to more than 25 transcription factors and were thus removed from the analysis.

Protein interactions were obtained from the *SIGNOR* [35] database (downloaded July 3, 2017). We deleted interactions annotated as “transcriptional regulation”, “guanine nucleotide exchange factor”, “transcriptional activation”, “post transcriptional regulation”, or “transcriptional repression”. Then, we deleted nodes that do not correspond to proteins, their complexes or families by retaining only nodes with *UniProt* [36] or *SIGNOR* ID.

Betweenness analysis was performed in R (3.4.0) using the *igraph* package (1.1.2). All shortest paths from TGF-β receptors to a given gene list were obtained using the igraph function *all_shortest_paths*. Then, the number of occurrences of each protein in these paths was calculated and used as a measure of graph betweenness. Finally, betweenness of cluster B was compared to betweenness of markers of other clusters.

**Transcription factor enrichment analysis**
To identify transcription factors with target genes enriched among IRCFs, we performed enrichment using the *Enrichr* [32] software and gene sets *TRANSFAC*_and_*JASPAR_PWMs* *ENCODE_TF_ChIP-seq_2015* *ENCODE*_and_*ChEA_Consensus_TFs*_from_*ChIP*-X *ChEA_2016*. p-values were adjusted using the R function *p.adjust* with method "BH". IRCFs with adjusted p-value < 0.05 were used in the enrichment.

**Data availability or Data resources**
Single-cell RNA-seq data was deposited at NBCI's Sequence Read Archive database under accession number SRAXXXX. RNA-seq data was deposited at NBCI's Sequence Read Archive database under accession number SRAXXXX. ATAC-seq data was deposited at NBCI's Sequence Read Archive database under accession number SRAXXXX. RNA-seq data was deposited at NBCI's XXXXX database under accession number XXXX (Cardiomyocytes).

**Supplementary Table 1- List of antibodies used in FACS and cell sorting**

| Name | Clone | Supplier | Dilution |
|---|---|---|---|
| CD11b-APC/Cy7 | M1/70 | BioLegend, 101225 | 1:100 |
| CD31 (PECAM)-APC | MEC 13.3 | BD Pharmingen, 551262 | 1:200 |
| CD31 (PECAM)-PE | MEC 13.3 | BD Pharmingen, 555027 | 1:200 |
| CD31-APC | 390 | BioLegend, 102409 | 1:200 |
| CD45-BV510 | 30-F11 | BioLegend, 103137 | 1:200 |
| CD45-PE | 30-F11 | eB 12-0451-81 | 1:200 |
| CD45-PerCP | 30-F11 | BD Pharmingen, 557235 | 1:200 |
| CD45-FITC | 30-F11 | BioLegend, 103107 | 1:100 |
| CD90.2-PE/Cy7 | 53-2.1 | eB 25-0902-82 | 1:100 |
| CD140a-PE/Cy7 | APA5 | eB25-1401-80 | 1:100 |
| CD146-PE/Cy7 | ME-9F1 | Biolegend 134714 | 1:100 |
| CD200-BV421 | OX-90 | BD Bioscience 565547 | 1:100 |
| Anti-feeder cells-APC | mEFSK4 | Miltenyi, 130-102-302 | 1:50 |
| Anti-CTHRC1 serum | pAb | Paygay et al, 2005 | 1:100 |
| Anti-rabbit IgG- PE | - | Jackson Inmunoresearch, 111-116-144 | - |

**Supplementary Table 2- List of primary and secondary antibodies used in Western Blot**

| Primary antibodies | Clone | Supplier | Dilution |
|---|---|---|---|
| Procollagen I | SP1.D8 | DSHB | 1:1000 |
| CTHRC1 | pAb | Maine Medical Center Research Institute, Vli08G09 | 1:1000 |
| β-Actin | pAb | Sigma-Aldrich, A2103 | 1:5000 |
| α-Tubulin | DM1A | Sigma-Aldrich, T9026 | 1:5000 |
| Phospho-Akt (Ser473) | pAb | Cell Signalling, 9271 | 1:1000 |
| Total Akt | pAb | Cell Signalling, 9272 | 1:1000 |
| Donkey HRP-conjugated anti-rabbit | pAb | GE Healthcare, NA934 | 1:10000 |
| Sheep HRP-conjugated anti-mouse | pAb | GE Healthcare, NA931 | 1:10000 |

**Supplementary Table 3- List of primary antibodies used in immunohistochemistry**

| Name | Clone | Supplier | Dilution |
|---|---|---|---|
| CD31 (PECAM) | MEC 13.3 | BD Pharmingen, 550274 | 1:100 |
| CD45-PE | 30-F11 | eB 12-0451-81 | 1:100 |
| CD200-BV421 | OX-90 | BD Bioscience, 565547 | 1:200 |
| Collagen-I | pAb | Rockland, 600-401-1035 | 1:200 |
| CTHRC1 | pAb | Maine Medical Center Research Institute (Vli55) | 1:50 |
| GFP | pAb | Abcam, ab13970 | 1:100 |

| | | | |
|---|---|---|---|
| Ki67 | pAb | Abcam, ab15580 | 1:50 |
| Sarcomeric α-Actin (αSA) | EA-53 | SIGMA, A7811 | 1:100 |
| Cardiac Troponin 1 (TnT) | 4C2 | GeneTex, GTX10231 | 1 ug/ml |
| Periostin (POSTN) | pAb | Abcam, ab79946 | 1:100 |

**Supplementary Table 4- List of secondary antibodies used in immunohistochemistry**

| **Name** | **Supplier** | **Dilution** |
|---|---|---|
| Anti-Chicken 488 | Jackson Inmunoresearch, 703-545-155 | 1:200 |
| Anti-Rabbit 488 | Invitrogen, A11008 | 1:100 |
| Anti-Rabbit 594 | Invitrogen, A21207 | 1:200 |
| Anti-Rat 594 | Invitrogen, A11007 | 1:200 |
| Anti-Mouse 546 | Invitrogen, A11003 | 1:100 |
| Anti-Mouse 647 | Invitrogen, A31571 | 1:200 |
| Anti-Rabbit 647 | Invitrogen, A31573 | 1:200 |
| Anti-Rat 647 | Invitrogen, A21247 | 1:200 |

**Supplementary Table 5- List of primers used in swine qPCR analysis**

| Name | Fwd | Rev |
|---|---|---|
| *HPRT1* | CCCAGCGTCGTGATTAGTGAT | TCCAGCAGGTCAGCAAAGAA |
| *RPL4* | GTTGGCATCGCAGAGTGAAC | CTCATTCGCTGAGAGGCA |
| *COL1α1* | CCTCAAGAGAAGGCTCACGA | TTGGTTGGGGTCAATCCAGT |
| *CTHRC1* | GGTCGGGATGGATTCAAA | TTAGGGCACTGTTGGAACGC |
| *POSTN* | TGTGCCAACCAATGATGCCT | ACAGGTATGTCTGCTGGGTAG |

### REFERENCES


1. Yata, Y., *et al.* DNase I-hypersensitive sites enhance alpha1(I) collagen gene expression in hepatic stellate cells. *Hepatology* **37**, 267-276 (2003).
2. Stohn, J.P., Perreault, N.G., Wang, Q., Liaw, L. & Lindner, V. Cthrc1, a novel circulating hormone regulating metabolism. *PLoS One* **7**, e47142 (2012).
3. Ruiz-Villalba, A., *et al.* Interacting resident epicardium-derived fibroblasts and recruited bone marrow cells form myocardial infarction scar. *J Am Coll Cardiol* **65**, 2057-2066 (2015).
4. Benavides-Vallve, C., *et al.* New strategies for echocardiographic evaluation of left ventricular function in a mouse model of long-term myocardial infarction. *PLoS One* **7**, e41691 (2012).
5. Doring, H.J. The isolated perfused heart according to Langendorff technique--function--application. *Physiol Bohemoslov* **39**, 481-504 (1990).
6. Jaitin, D.A., *et al.* Massively parallel single-cell RNA-seq for marker-free decomposition of tissues into cell types. *Science* **343**, 776-779 (2014).
7. Lavin, Y., *et al.* Innate Immune Landscape in Early Lung Adenocarcinoma by Paired Single-Cell Analyses. *Cell* **169**, 750-765 e717 (2017).
8. Bagnoli, J.W., *et al.* Sensitive and powerful single-cell RNA sequencing using mcSCRB-seq. *Nature communications* **9**, 2937 (2018).
9. Lara-Astiaso, D., *et al.* Immunogenetics. Chromatin state dynamics during blood formation. *Science* **345**, 943-949 (2014).
10. Corces, M.R., *et al.* Lineage-specific and single-cell chromatin accessibility charts human hematopoiesis and leukemia evolution. *Nat Genet* **48**, 1193-1203 (2016).
11. Buenrostro, J.D., Giresi, P.G., Zaba, L.C., Chang, H.Y. & Greenleaf, W.J. Transposition of native chromatin for fast and sensitive epigenomic profiling of open chromatin, DNA-binding proteins and nucleosome position. *Nat Methods* **10**, 1213-1218 (2013).
12. Jain, K., Verma, P.J. & Liu, J. Isolation and handling of mouse embryonic fibroblasts. *Methods Mol Biol* **1194**, 247-252 (2014).
13. Ruiz-Villalba, A., van Pelt-Verkuil, E., Gunst, Q.D., Ruijter, J.M. & van den Hoff, M.J. Amplification of nonspecific products in quantitative polymerase chain reactions (qPCR). *Biomol Detect Quantif* **14**, 7-18 (2017).
14. Ruijter, J.M., *et al.* Amplification efficiency: linking baseline and bias in the analysis of quantitative PCR data. *Nucleic Acids Res* **37**, e45 (2009).
15. Ruijter, J.M., Ruiz Villalba, A., Hellemans, J., Untergasser, A. & van den Hoff, M.J. Removal of between-run variation in a multi-plate qPCR experiment. *Biomol Detect Quantif* **5**, 10-14 (2015).
16. Ruiz-Villalba, A., *et al.* Reference genes for gene expression studies in the mouse heart. *Sci Rep* **7**, 24 (2017).
17. Nygard, A.B., Jorgensen, C.B., Cirera, S. & Fredholm, M. Selection of reference genes for gene expression studies in pig tissues using SYBR green qPCR. *BMC Mol Biol* **8**, 67 (2007).
18. Hellemans, J., Mortier, G., De Paepe, A., Speleman, F. & Vandesompele, J. qBase relative quantification framework and software for management and automated analysis of real-time quantitative PCR data. *Genome Biol* **8**, R19 (2007).
19. Villani, A.C., *et al.* Single-cell RNA-seq reveals new types of human blood dendritic cells, monocytes, and progenitors. *Science* **356**(2017).
20. Butler, A., Hoffman, P., Smibert, P., Papalexi, E. & Satija, R. Integrating single-cell transcriptomic data across different conditions, technologies, and species. *Nat Biotechnol* **36**, 411-420 (2018).
21. Li, W., Cerise, J.E., Yang, Y. & Han, H. Application of t-SNE to human genetic data. *J Bioinform Comput Biol* **15**, 1750017 (2017).

22. Dobin, A., *et al.* STAR: ultrafast universal RNA-seq aligner. *Bioinformatics* **29**, 15-21 (2013).
23. Liao, Y., Smyth, G.K. & Shi, W. The R package Rsubread is easier, faster, cheaper and better for alignment and quantification of RNA sequencing reads. *Nucleic Acids Res* **47**, e47 (2019).
24. Robinson, M.D., McCarthy, D.J. & Smyth, G.K. edgeR: a Bioconductor package for differential expression analysis of digital gene expression data. *Bioinformatics* **26**, 139-140 (2010).
25. Love, M.I., Huber, W. & Anders, S. Moderated estimation of fold change and dispersion for RNA-seq data with DESeq2. *Genome Biol* **15**, 550 (2014).
26. Langmead, B., Trapnell, C., Pop, M. & Salzberg, S.L. Ultrafast and memory-efficient alignment of short DNA sequences to the human genome. *Genome Biol* **10**, R25 (2009).
27. Zhang, Y., *et al.* Model-based analysis of ChIP-Seq (MACS). *Genome Biol* **9**, R137 (2008).
28. Zhu, L.J., *et al.* ChIPpeakAnno: a Bioconductor package to annotate ChIP-seq and ChIP-chip data. *BMC bioinformatics* **11**, 237 (2010).
29. Yu, G., Wang, L.G., Han, Y. & He, Q.Y. clusterProfiler: an R package for comparing biological themes among gene clusters. *OMICS* **16**, 284-287 (2012).
30. Fabregat, A., *et al.* The Reactome Pathway Knowledgebase. *Nucleic Acids Res* **46**, D649-D655 (2018).
31. Ferreira, J.A. The Benjamini-Hochberg method in the case of discrete test statistics. *Int J Biostat* **3**, Article 11 (2007).
32. Kuleshov, M.V., *et al.* Enrichr: a comprehensive gene set enrichment analysis web server 2016 update. *Nucleic Acids Res* **44**, W90-97 (2016).
33. Rouillard, A.D., *et al.* The harmonizome: a collection of processed datasets gathered to serve and mine knowledge about genes and proteins. *Database (Oxford)* **2016**(2016).
34. Han, H., *et al.* TRRUST v2: an expanded reference database of human and mouse transcriptional regulatory interactions. *Nucleic Acids Res* **46**, D380-D386 (2018).
35. Perfetto, L., *et al.* SIGNOR: a database of causal relationships between biological entities. *Nucleic Acids Res* **44**, D548-554 (2016).
36. The UniProt, C. UniProt: the universal protein knowledgebase. *Nucleic Acids Res* **45**, D158-D169 (2017).
