## Supplementary figures and images for "Single-cell RNA-seq analysis reveals the crucial role of Collagen Triplex Helix Repeat Containing 1 (CTHRC1) cardiac fibroblasts for ventricular remodeling after myocardial infarction"

### Supplementary Figure 1

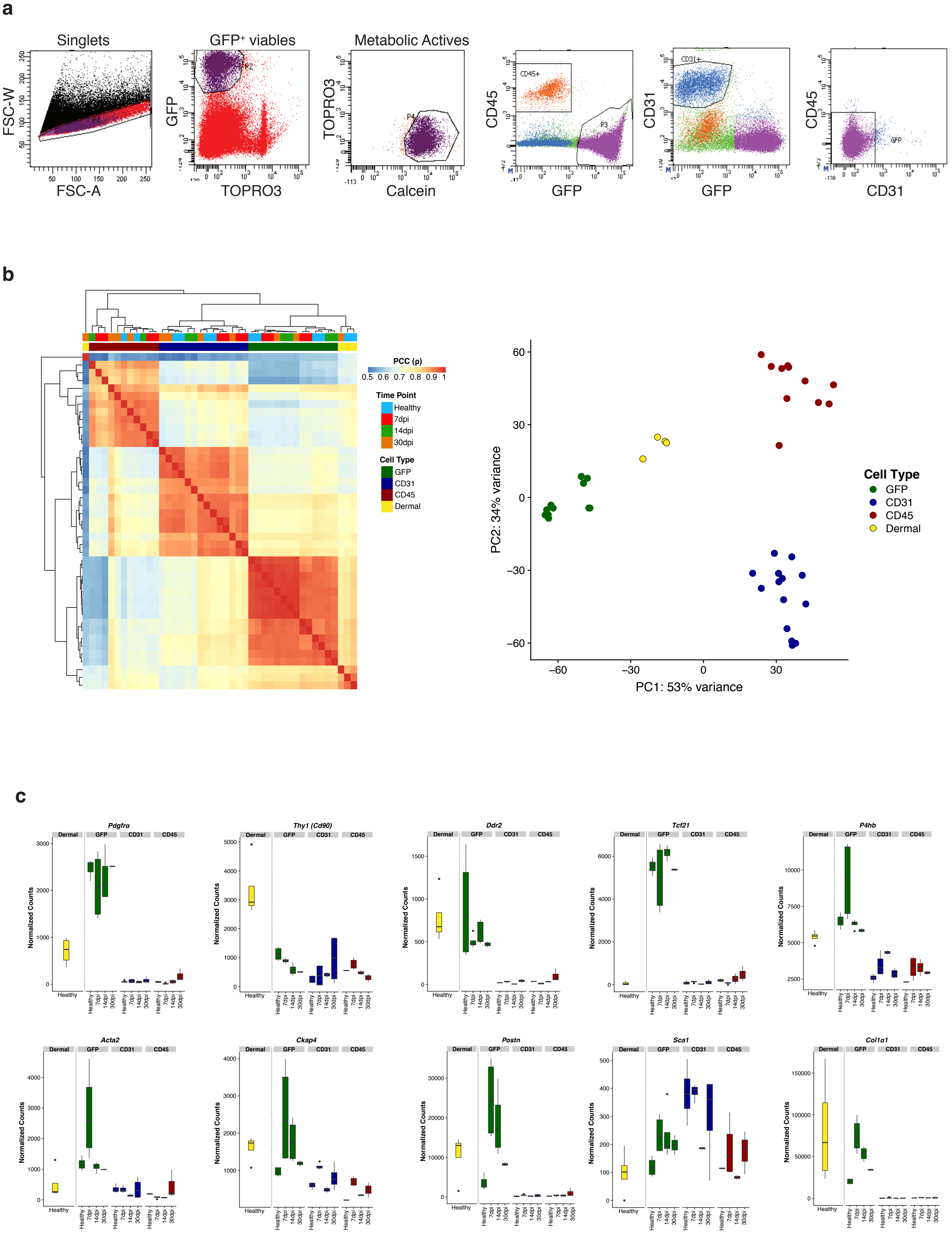

### Supplementary Figure 2

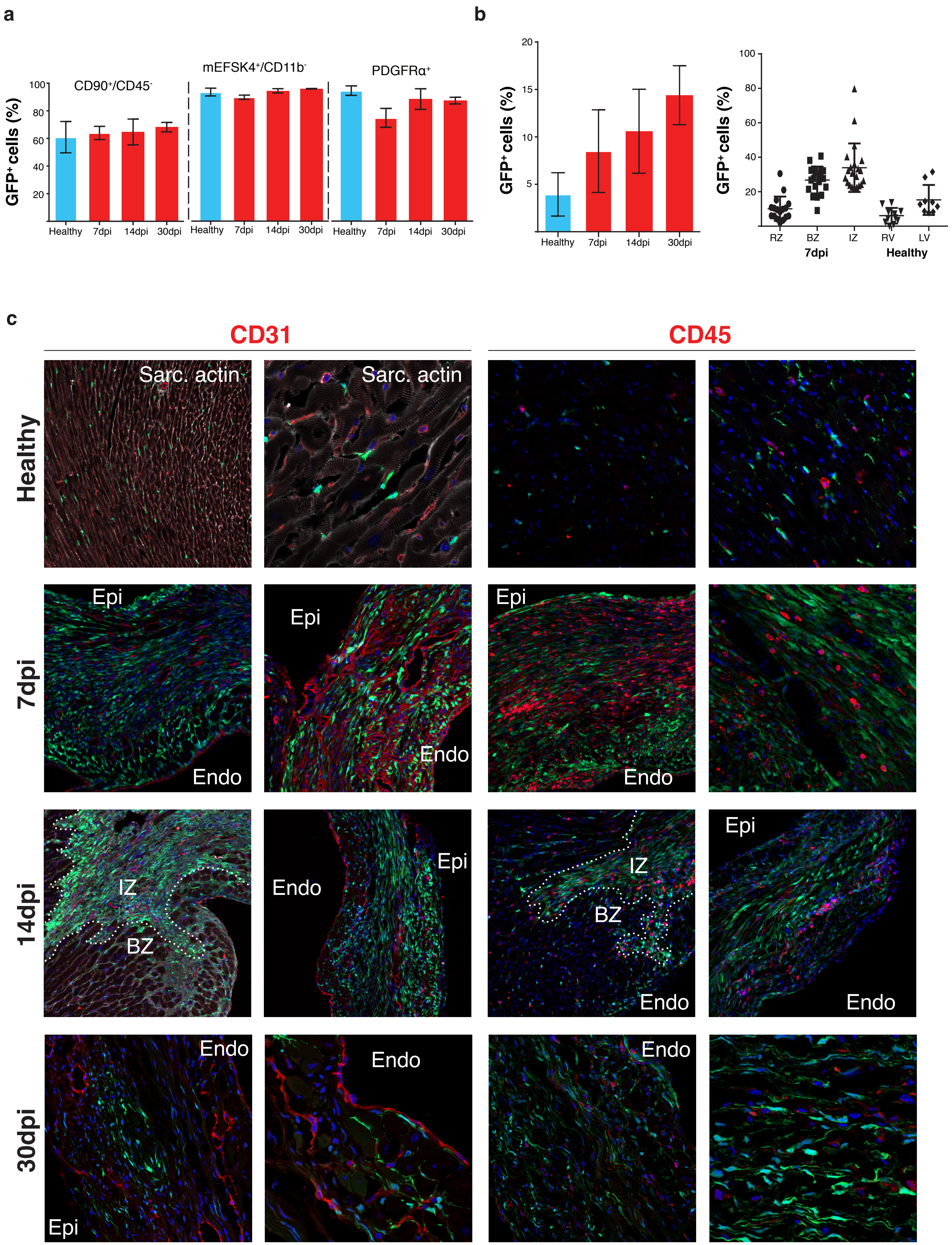

### Supplementary Figure 3

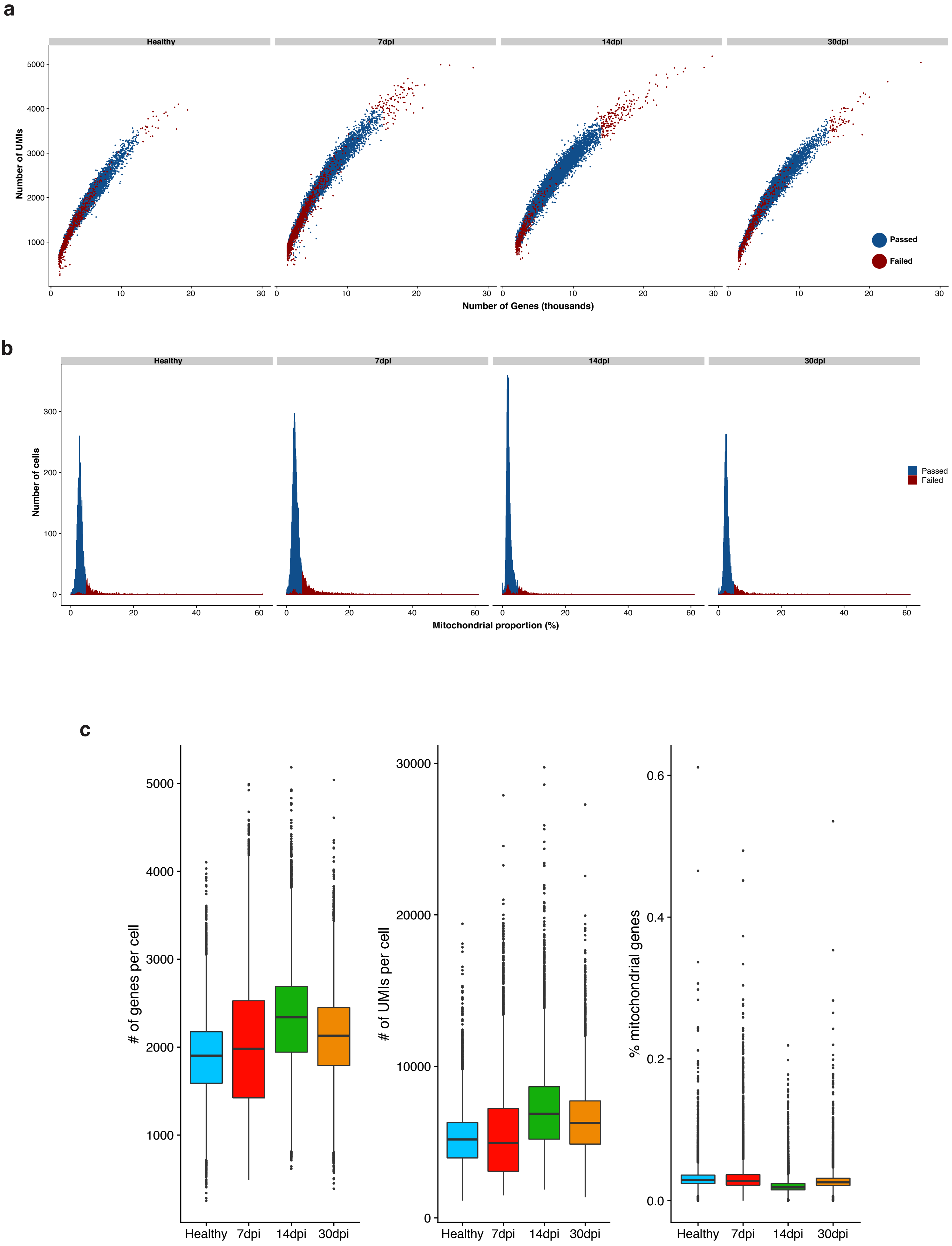

### Supplementary Figure 4

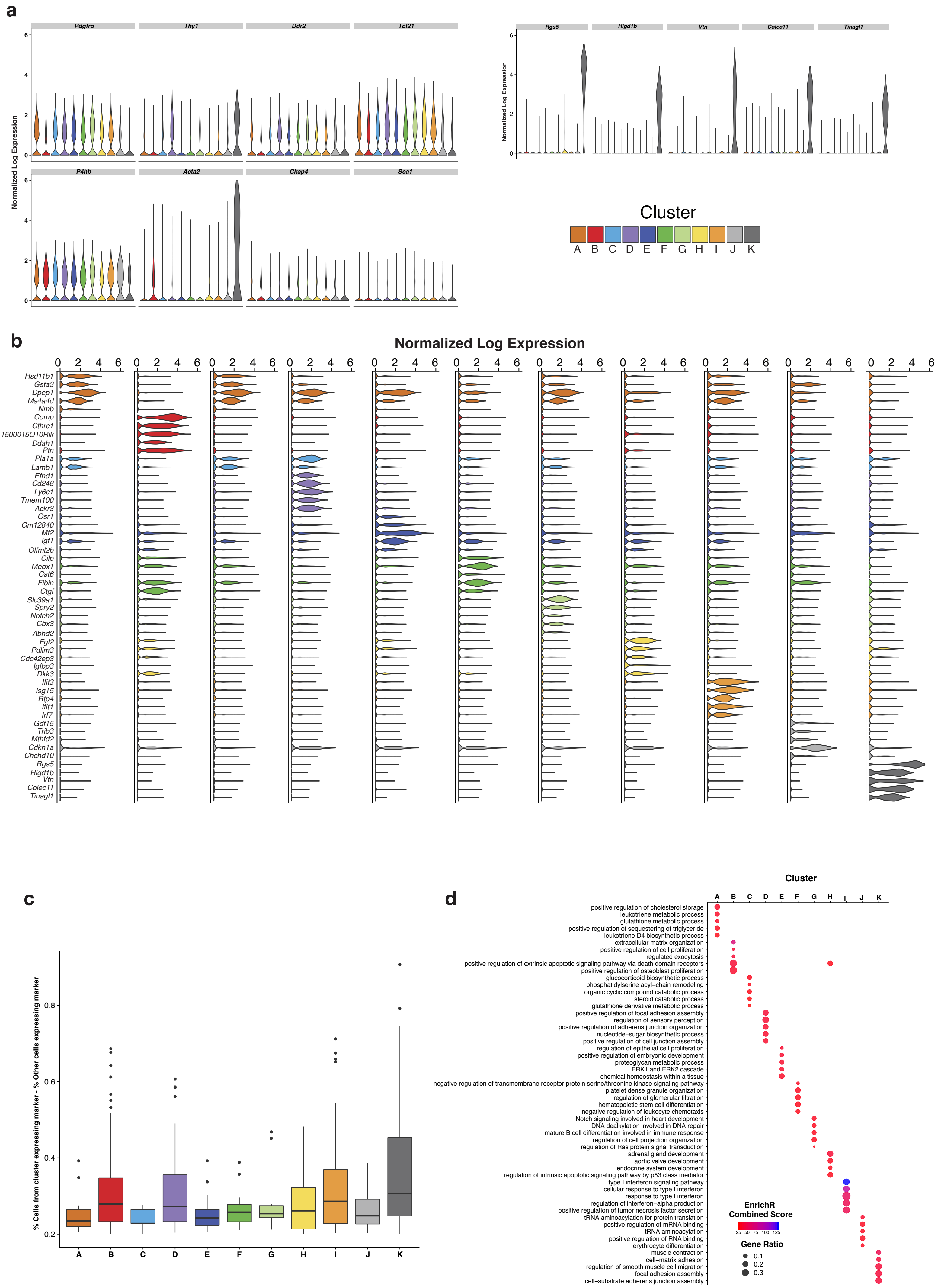

### Supplementary Figure 5

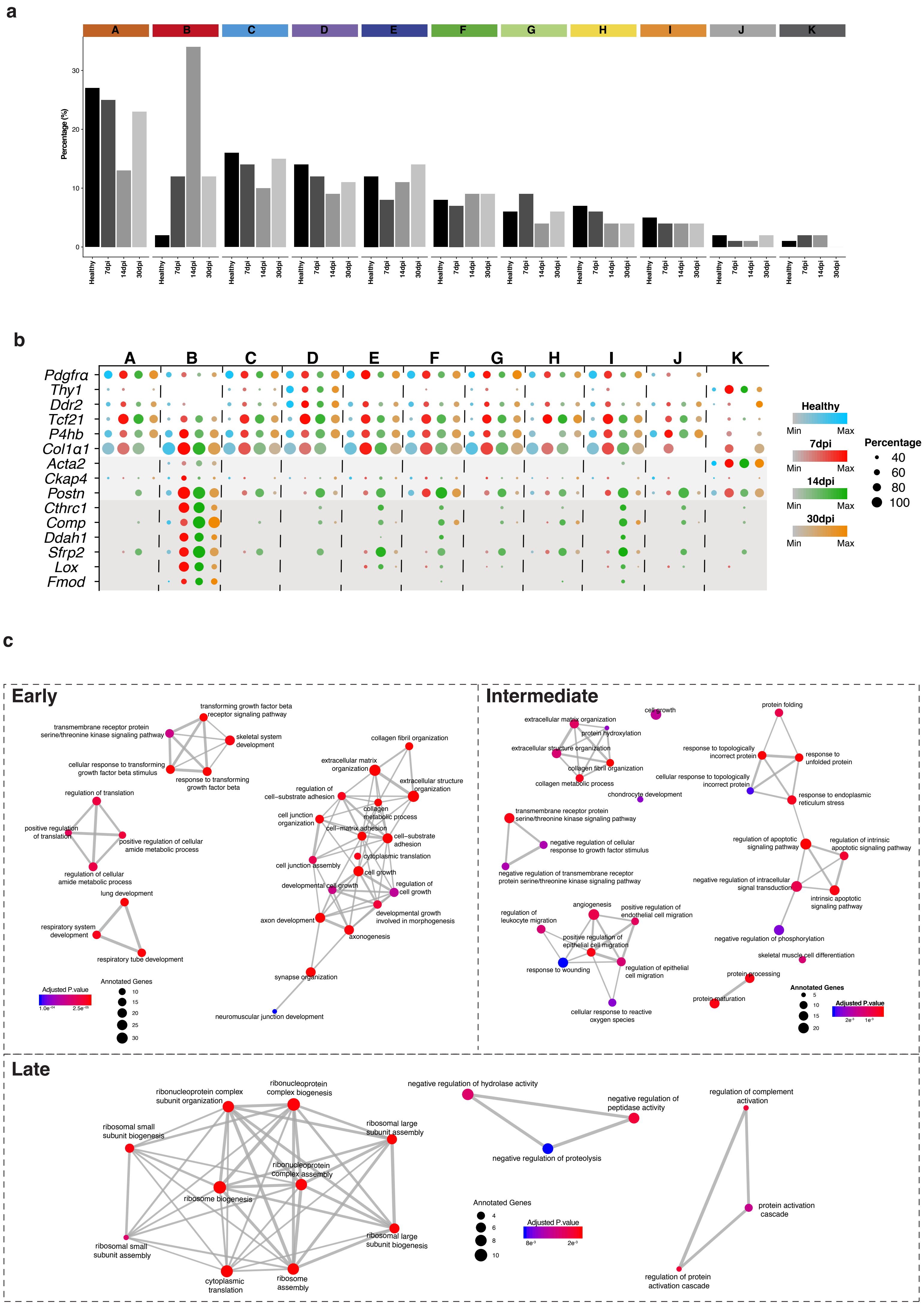

### Supplementary Figure 6

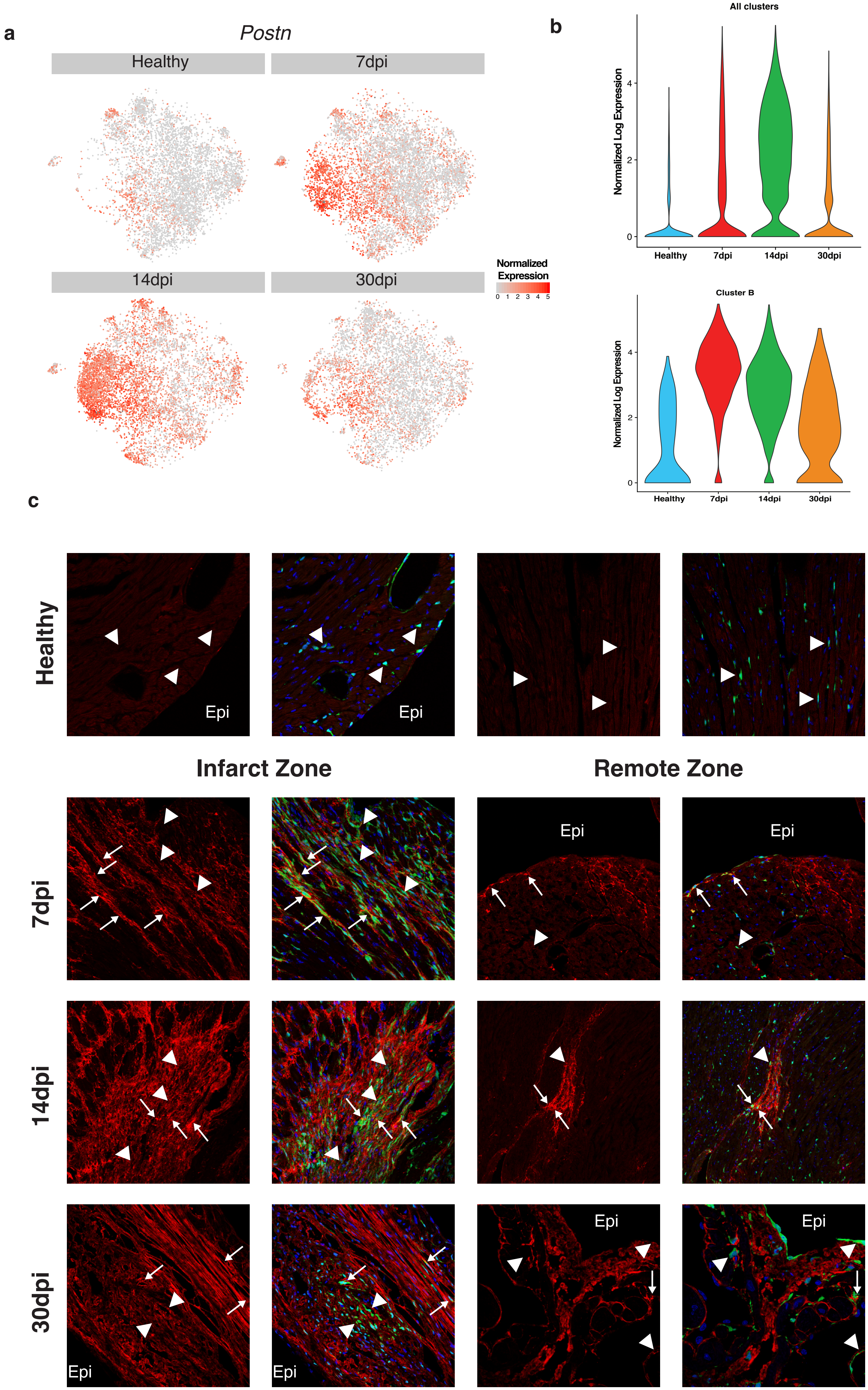

### Supplementary Figure 7

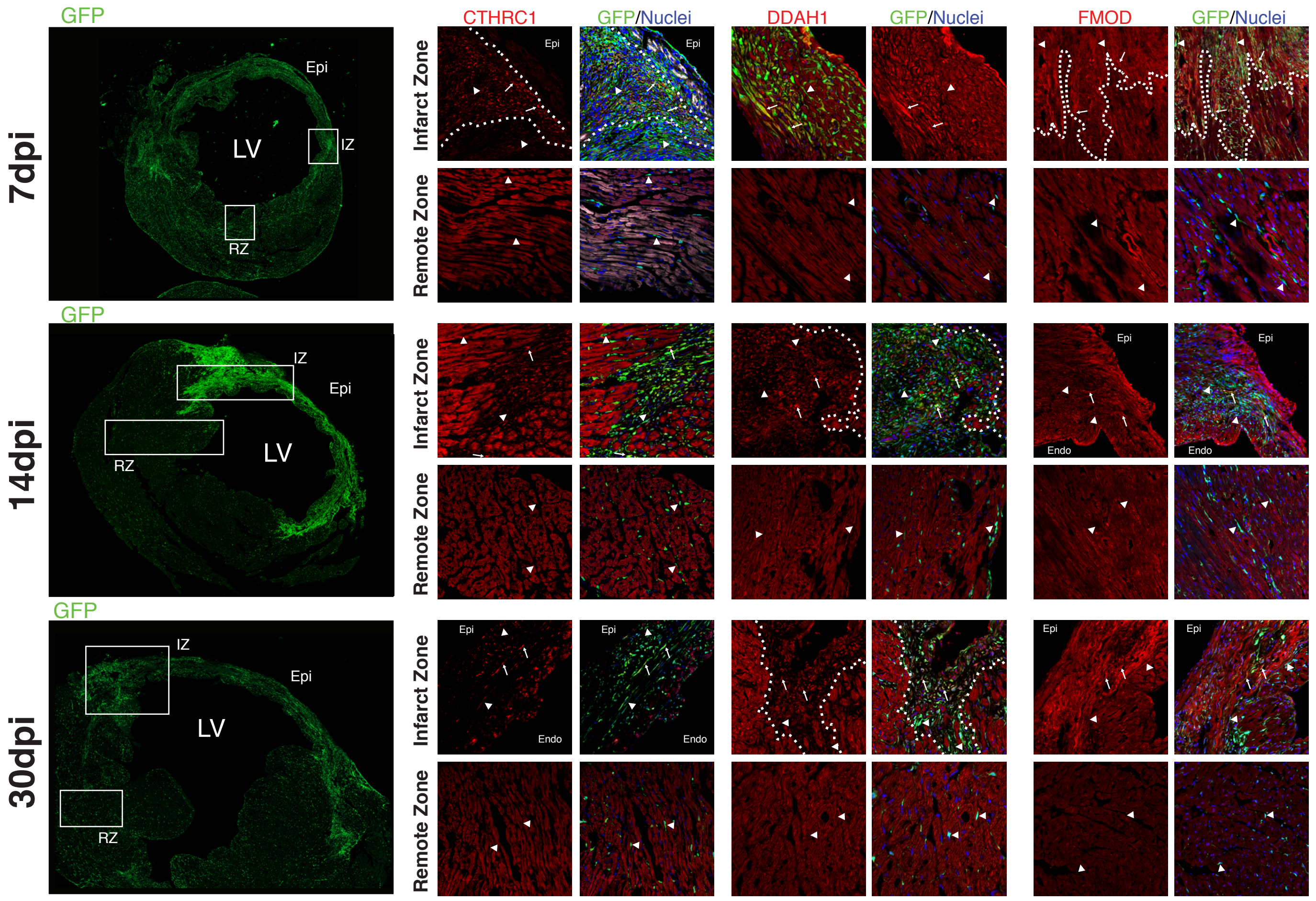

### Supplementary Figure 8

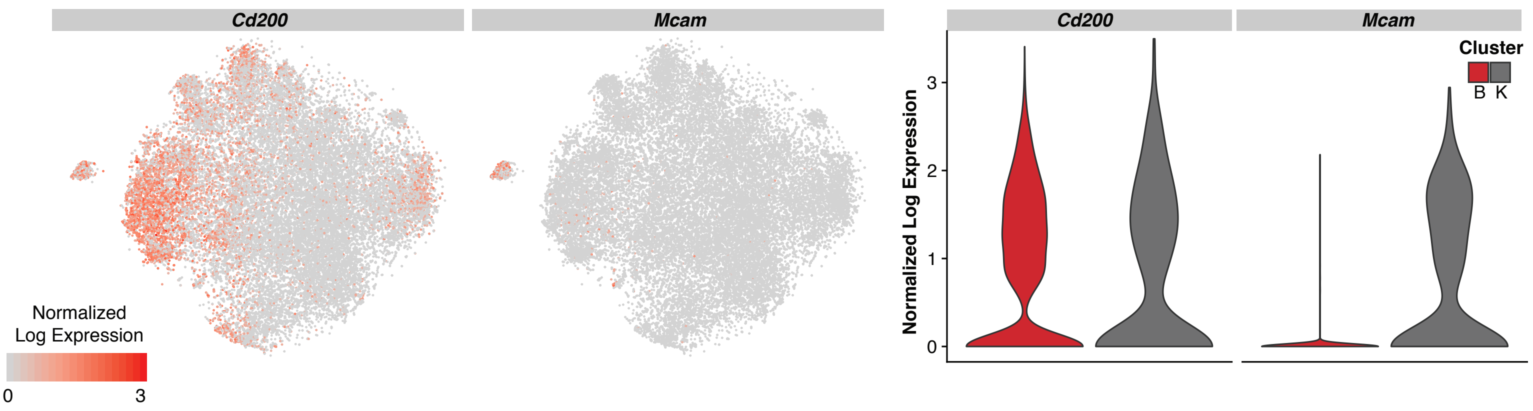

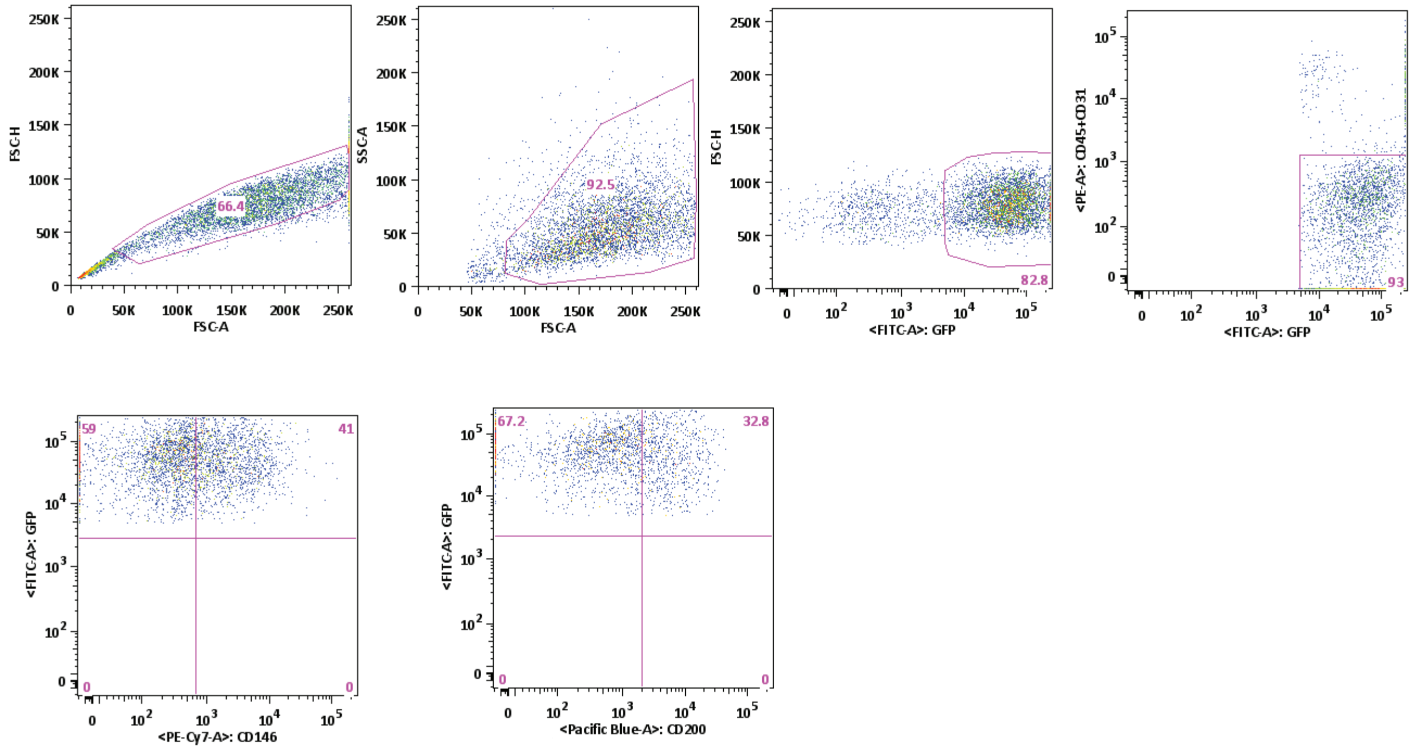

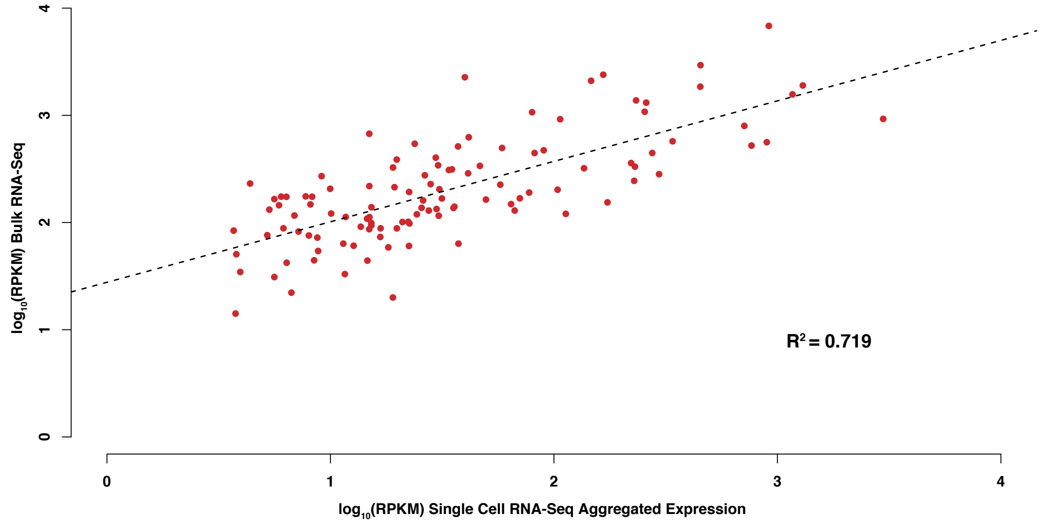

### Supplementary Figure 9

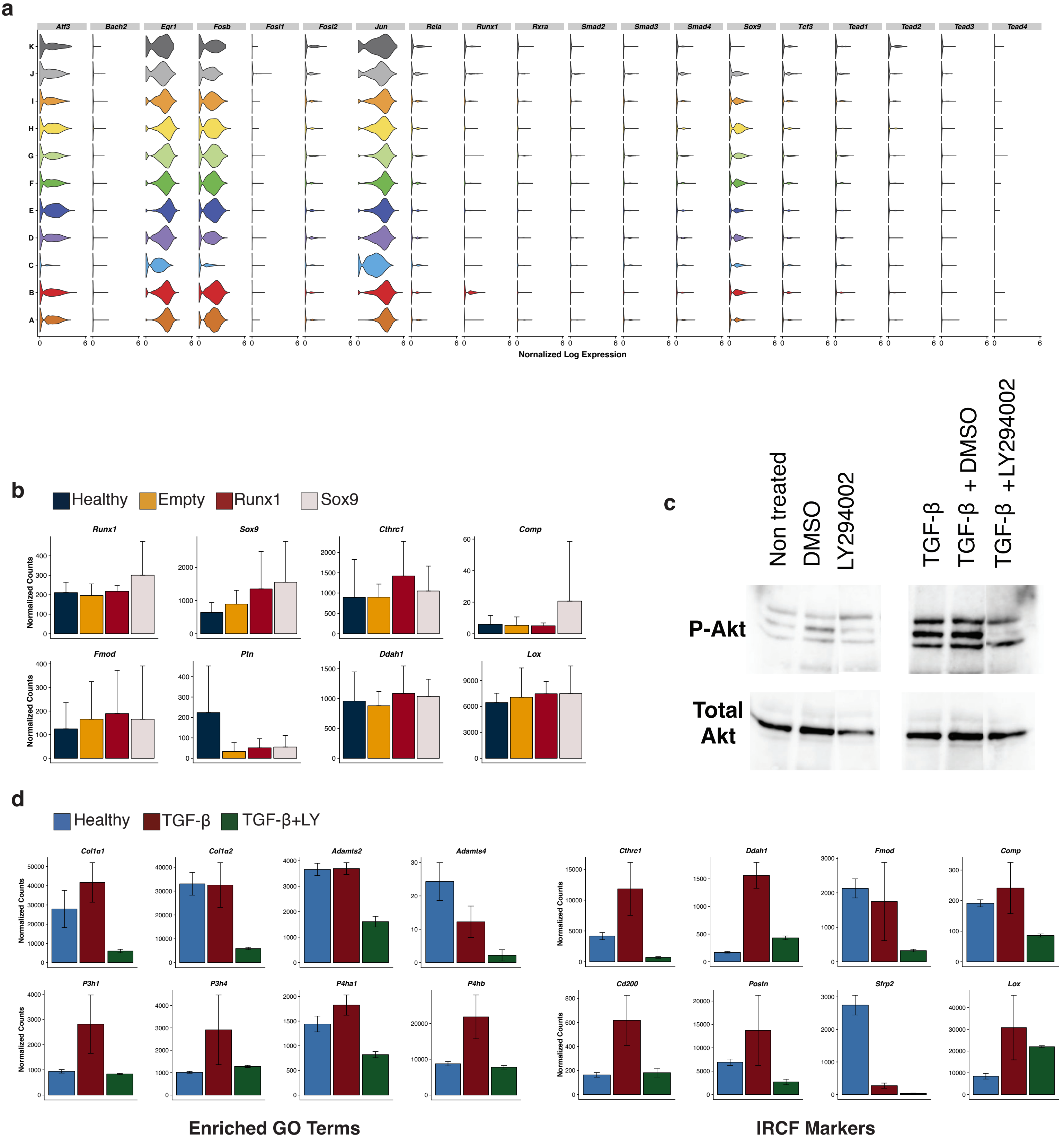

### Supplementary Figure 10

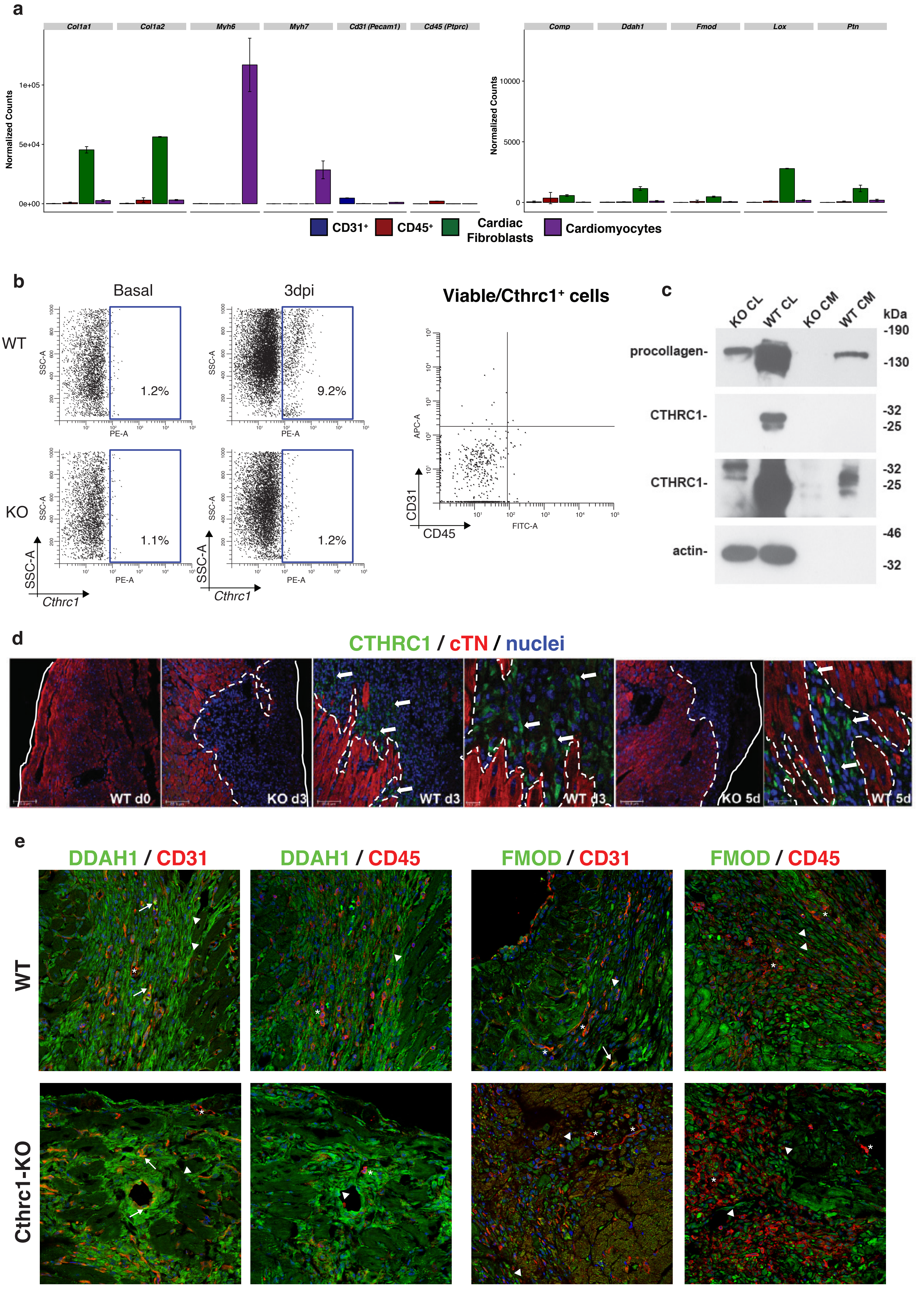
